# The human metabolite–protein interactome reveals a global layer of cellular coordination

**DOI:** 10.64898/2026.08.28.747909

**Authors:** Jeffrey Skolnick, Bharath Srinivasan

**Affiliations:** Parker H. Petit Center for AI-Driven Health Innovation, Georgia Institute of Technology, 950 Atlantic Dr NW, Atlanta, GA 30332, USA; School of Pharmacy and Life Sciences, Robert Gordon University, Garthdee House, Garthdee Rd, Aberdeen AB10 7AQ, UK; Department of Chemistry, Stony Brook University, Stony Brook, NY 11794-3400, USA; Center for the Advanced Study of Drug Action, Stony Brook University, Stony Brook, NY 11794-3400, USA; Cancer Research Horizons, Cancer Research UK, 2 Redman Place, London E20 1JQ, UK

## Abstract

Metabolites are substrates, products, cofactors, and regulators, but protein–protein interaction networks do not represent their potential to organize proteins across conventional pathway boundaries. Using the LIGMAP virtual-screening algorithm, we mapped 308 human metabolite codes to pockets in monomers, dimer interfaces, and non-interface sites in dimers and represented attractive or repulsive COLIG states involving pairs of metabolites in the same pocket. On a fixed cohort of 3,938 proteins, the mean coverage of 104 strict non-enzyme pathways was 44.6% for LIGMAP, 70.1% for STRING, and 82.5% for STRING+LIGMAP; the union placed 86.3% of eligible proteins in the largest connected component and 95.1% in the two largest components. STRING+LIGMAP protein coverage was 88.0% for 68 enzyme-only pathways and 88.9% for 1,283 mixed pathways. In pathway-held-out, degree-matched prediction, adding LIGMAP to degree plus STRING increased the mean area under the precision–recall curve from 0.651 to 0.660 (paired P = 0.024); adding BioLiP2 increased it to 0.663 (paired P = 0.005). Ancient-only and non-ancient-only subnetworks were each globally connected; ancient features were denser, whereas non-ancient features covered more proteins and pathways. At the full 5,426-protein scale, retaining only features assigned to 2–100 proteins recovered 698 of 1,691 strict non-enzyme reference edges (41.3%) and exceeded both protein-label and exact bipartite degree-preserving nulls. Uncapped recovery approached saturation and lost identity-selective enrichment. Experimentally established metabolite-dependent complexes validate the local mechanism independently of LIGMAP; LIGMAP fully recovered two of seven stringent direct mechanisms and all three broader serial axes examined. Our findings reveal a global metabolite-mediated architecture with the capacity to coordinate proteins across otherwise distinct cellular systems.

## Introduction

Protein function is commonly organized into reaction pathways, physical interaction networks, and spatially assembled metabolons (1–8). Yet metabolites also bind non-enzymes, allosteric sites, and protein interfaces, and proteome-scale experiments reveal extensive protein–metabolite communication (9–17). A shared metabolite can therefore couple proteins that may not form a stable protein–protein interaction and may belong to different curated pathways.

The Entabolon hypothesis (18) generalizes this observation: overlapping metabolite-binding relationships could form a latent, context-gated protein network whose expressed state depends on protein association, abundance, compartment, affinity, competition within and between binding sites, post-translational state, and timing. Proteome-scale studies have established that protein–metabolite interactions are widespread and can regulate individual complexes, enzymes, and pathways (9–17). What remains unresolved is whether these local mechanisms collectively generate a global organizational layer across the human proteome. Addressing that question requires sufficiently comprehensive data for the combined human proteome and metabolome.

Because proteome-wide protein–metabolite binding information remains incomplete, computational predictions can help fill this gap. LIGMAP predicts metabolite binding by aligning CAVITATOR-defined pockets with APoc (18–21), and COLIG analysis extends the representation to attractive or repulsive metabolite pairs occupying the same pocket or interface (22).

As shown schematically in Figure 1, we first ask a narrow quantitative question: how much curated pathway organization is recovered by LIGMAP (18), STRING protein–protein interactions (7), BioLiP2 experimental protein–ligand contacts (23), and their unions? Reactome (8) supplies the primary pathway benchmark, and WikiPathways (24) tests whether the conclusions persist under a community-curated alternative. We then examine protein coverage, largest- and two-largest-component coverage, and enzyme-assisted connection of non-enzyme cores. Only after these benchmarks do we ask what ancient metabolites contribute, why dense metabolite networks become globally connected, and which experimental systems support the Entabolon mechanism. This progression tests whether metabolite-mediated organization is merely a collection of local, idiosyncratic effects or a general property of the protein–metabolite system.

**Figure 1.**
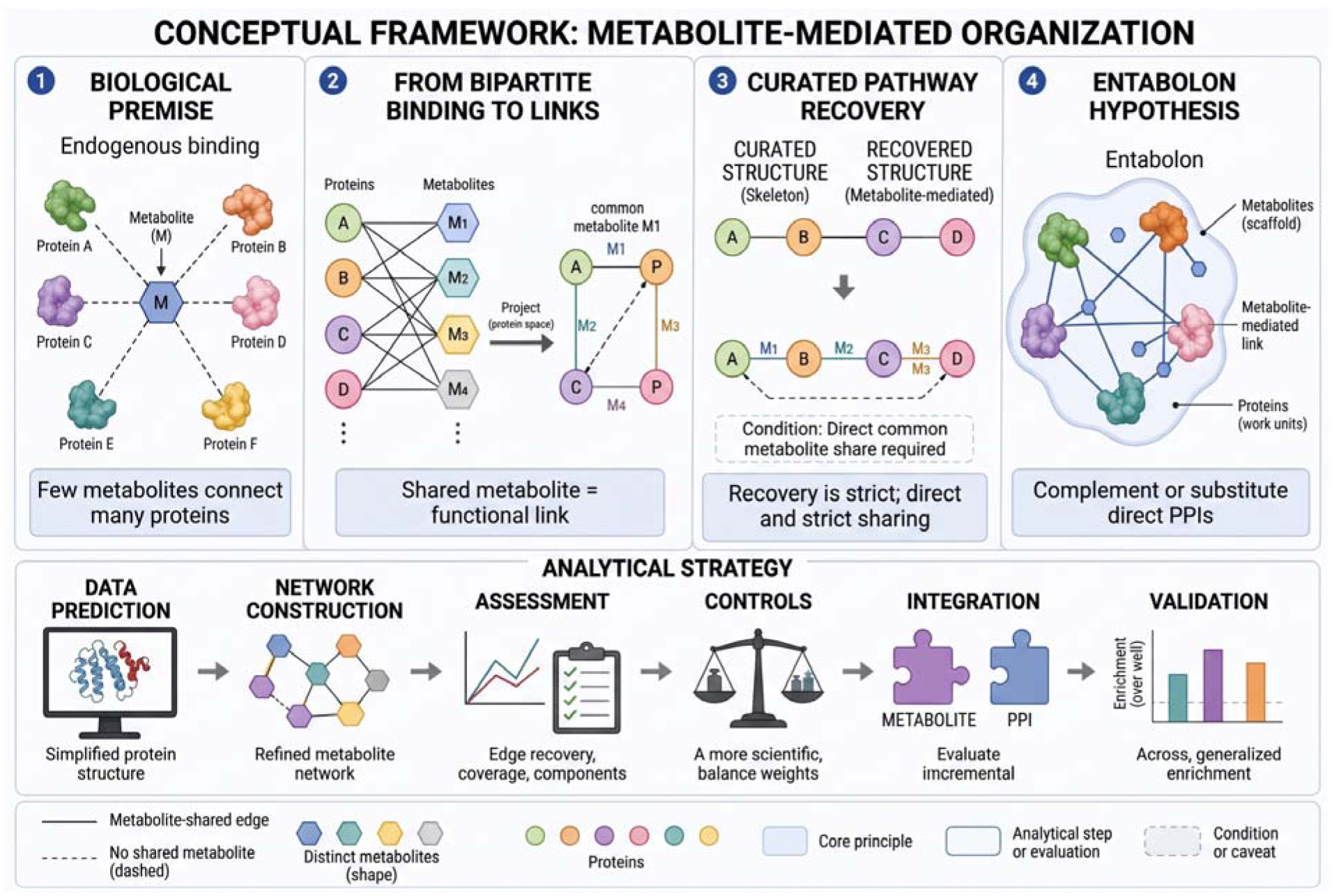
Integrated conceptual framework for metabolite-enabled pathway and global protein organization. The analysis begins with 308 metabolite codes mapped by LIGMAP to CAVITATOR-defined, APoc-aligned pockets in standalone monomers, protein–protein interfaces of structurally observed dimers, and non-interface sites in dimer subunits. Single-metabolite assignments and attractive or repulsive COLIG states (22) are projected from a protein– metabolite bipartite representation into protein–protein coupling edges. COLIGs are pairs of ligands predicted to occupy the same protein pocket. These metabolite-mediated edges complement high-confidence physical STRING interactions and are evaluated by recovery of non-enzyme, enzyme-only, and mixed pathways using protein coverage and largest- and two-largest-component statistics. Pathway-held-out prediction, structural-layer ablation, feature-degree-cap sensitivity, experimental BioLiP2 contacts, and degree-preserving null networks distinguish incremental biological placement from the general connectivity produced by a dense incidence architecture. Ancient and non-ancient metabolites provide differently distributed but convergent contributions to the connected network. The resulting graph is interpreted as a latent architecture rather than a constitutively active cell-wide complex: metabolite abundance, compartment, affinity, competition, protein state, and timing select context-expressed Entabolon subnetworks. Experimentally established metabolite-dependent complex stabilization, disruption, and downstream cellular responses validate local mechanisms implied by the Entabolon hypothesis; the MTAP–MTA– PRMT5 axis, including predicted repulsive MTA|SAM occupancy, provides a causal and reversible example. Thus, conventional pathways serve as a quantitative benchmark, whereas collective, context-dependent metabolite-enabled organization is the broader implication.

## Results

### Pathway reconstruction by LIGMAP, STRING, BioLiP, and their unions

As shown in Table 1, STRING alone was strongest for strict non-enzyme (104) and mixed (1,283) pathways, whereas LIGMAP alone was strongest for enzyme-only (68) pathways. One-component reconstruction means that all eligible pathway proteins form one connected component; the largest-component percentage instead reports the mean fraction of eligible proteins in the largest component. The union consistently improved reconstruction, demonstrating that metabolite-mediated edges are not redundant with high-confidence protein– protein interactions. BioLiP2 added modest coverage because experimentally resolved human metabolite-bound structures remain sparse, but it provided an orthogonal structural evidence layer and modestly improved enzyme-only and mixed-pathway reconstruction.

**Table 1.** Fixed-cohort pathway reconstruction and component coverage.

|  | Pathway class | One component | <=2 components | Protein coverage | Largest component<sup>a</sup> | Top two components<sup>a</sup> |
| --- | --- | --- | --- | --- | --- | --- |
| STRING | Non-enzyme | 53/104 (51.0%) | 76/104 (73.1%) | 70.1% | 74.9% | 89.5% |
| STRING | Enzyme-only | 25/68 (36.8%) | 39/68 (57.4%) | 67.1% | 65.2% | 82.8% |
| STRING | Mixed | 418/1283 (32.6%) | 673/1283 (52.5%) | 77.1% | 71.0% | 84.9% |
| LIGMAP | Non-enzyme | 29/104 (27.9%) | 69/104 (66.3%) | 44.6% | 62.4% | 86.9% |
| LIGMAP | Enzyme-only | 23/68 (33.8%) | 50/68 (73.5%) | 61.9% | 71.1% | 91.0% |
| LIGMAP | Mixed | 170/1283 (13.3%) | 459/1283 (35.8%) | 61.3% | 64.6% | 79.4% |
| STRING+LIGMAP | Non-enzyme | 73/104 (70.2%) | 87/104 (83.7%) | 82.5% | 86.3% | 95.1% |
| STRING+LIGMAP | Enzyme-only | 44/68 (64.7%) | 59/68 (86.8%) | 88.0% | 86.1% | 96.4% |
| STRING+LIGMAP | Mixed | 767/1283 (59.8%) | 1068/1283 (83.2%) | 88.9% | 88.8% | 96.4% |
| LIGMAP+BioLiP2 | Non-enzyme | 30/104 (28.8%) | 72/104 (69.2%) | 48.0% | 64.3% | 88.5% |
| LIGMAP+BioLiP2 | Enzyme-only | 27/68 (39.7%) | 52/68 (76.5%) | 66.8% | 73.7% | 92.3% |
| LIGMAP+BioLiP2 | Mixed | 187/1283 (14.6%) | 497/1283 (38.7%) | 64.9% | 66.8% | 81.8% |
| STRING+BioLiP2+LIGMAP | Non-enzyme | 73/104 (70.2%) | 88/104 (84.6%) | 82.6% | 86.5% | 95.2% |
| STRING+BioLiP2+LIGMAP | Enzyme-only | 46/68 (67.6%) | 59/68 (86.8%) | 89.0% | 87.1% | 96.6% |
| STRING+BioLiP2+LIGMAP | Mixed | 778/1283 (60.6%) | 1078/1283 (84.0%) | 89.3% | 89.1% | 96.6% |

<sup>a</sup>Fraction of proteins in the largest and first and second largest connected component of the pathway graph.

For each pathway, the network was restricted to the fixed set of eligible pathway proteins, including isolated proteins, and the resulting connected components were ranked by size. Largest-component coverage was defined as the number of proteins in the largest connected component divided by the total number of eligible pathway proteins. Top-two-component coverage was defined analogously as the combined number of proteins in the two largest connected components divided by the total eligible pathway-protein count. Values reported for each pathway class are means across all evaluable pathways. Thus, top-two-component coverage measures whether most pathway proteins can be represented by no more than two dominant network clusters and should not be confused with principal-component analysis.

STRING reconstructed more pathways than LIGMAP when each was used alone, but STRING+LIGMAP exceeded either individual method. Adding BioLiP2 to LIGMAP produced a smaller improvement, indicating that LIGMAP supplies most of the metabolite-associated network information in this comparison. Importantly, the two methods contributed partially overlapping but distinct edges: their union placed more than 95% of eligible proteins in the two largest components for all three Reactome pathway classes.

### Enzyme-assisted reconstruction of non-en**zyme cores**

Dividing pathways into enzyme-only, non-enzyme, and mixed classes is analytically useful but biologically artificial. Pathways dominated by non-enzymes provide a more stringent test of the Entabolon hypothesis because direct recovery of enzyme-containing pathways can partly reflect canonical substrate and product relationships. We therefore decomposed mixed pathways into their eligible non-enzyme core, the core plus at most one pathway enzyme, and the core plus all eligible pathway enzymes. For the one-enzyme sensitivity analysis, the enzyme maximizing the largest-component fraction was selected within each model; this is an upper bound on what one enzyme can contribute, not a prospective selection rule. STRING+LIGMAP increased mean non-enzyme-core coverage from 76.7% for the core alone to 90.3% with one enzyme and 91.7% with all eligible pathway enzymes. The corresponding top-two-component fractions were 93.3%, 97.0%, and 96.5%. Thus, a single pathway enzyme captured much of the attainable connectivity, and Tables 1 and 2 show that shared-metabolite and protein–protein interaction edges jointly recover substantial pathway organization.

**Table 2.** Enzyme-assisted connectivity of non-enzyme cores in mixed pathways.

| Network | Target | Number of Pathways | Protein coverage | Largest component<sup>a</sup> | Top two components<sup>a</sup> | One component<sup>b</sup> |
| --- | --- | --- | --- | --- | --- | --- |
| STRING | Core only | 1052 | 68.5% | 66.7% | 85.7% | 279 |
| STRING | Core +1 enzyme | 1052 | 80.3% | 75.5% | 88.9% | 397 |
| STRING | Core + all enzymes | 1052 | 80.6% | 71.8% | 84.4% | 329 |
| LIGMAP | Core only | 1052 | 40.4% | 51.5% | 73.2% | 75 |
| LIGMAP | Core +1 enzyme | 1052 | 59.7% | 62.3% | 77.7% | 149 |
| LIGMAP | Core + all enzymes | 1052 | 64.7% | 64.9% | 77.2% | 121 |
| STRING+LIGMAP | Core only | 1052 | 76.7% | 78.9% | 93.3% | 437 |
| STRING+LIGMAP | Core +1 enzyme | 1052 | 90.3% | 89.8% | 97.0% | 663 |
| STRING+LIGMAP | Core + all enzymes | 1052 | 91.7% | 90.3% | 96.5% | 635 |
| LIGMAP+BioLiP2 | Core only | 1052 | 44.6% | 53.5% | 75.5% | 87 |
| LIGMAP+BioLiP2 | Core +1 enzyme | 1052 | 63.3% | 64.4% | 80.1% | 159 |
| LIGMAP+BioLiP2 | Core + all enzymes | 1052 | 68.6% | 67.4% | 79.9% | 134 |
| STRING+BioLiP2+LIGMAP | Core only | 1052 | 77.8% | 80.1% | 94.3% | 446 |
| STRING+BioLiP2+LIGMAP | Core + 1 enzyme | 1052 | 90.5% | 89.9% | 97.1% | 666 |
| STRING+BioLiP2+LIGMAP | Core + all enzymes | 1052 | 92.0% | 90.6% | 96.8% | 644 |

<sup>a</sup>Fraction of proteins in the largest and first and second largest connected component of the pathway graph.
<sup>b</sup>Numbers of pathways for which all eligible pathway proteins form one connected component or no more than two connected components

### Comparison to WikiPathways

Reactome represents only one possible partition of proteins into pathways. We therefore repeated the analysis using the July 10, 2026 human WikiPathways collection (24). As shown in Table 3, the principal results persisted under the same fixed cohort, networks, cap, and component definitions. Among 110 strict non-enzyme pathways, mean protein coverage was 52.6% for STRING, 45.4% for LIGMAP, and 69.9% for STRING+LIGMAP; the union placed 74.9% and 92.1% of eligible proteins in the largest and two largest components, respectively. In 808 mixed pathways, LIGMAP alone exceeded STRING (77.3% versus 59.0%), while their union reached 88.7% coverage and the full union reached 89.9%. The smaller set of 17 enzyme-only pathways was weakly reconstructed by STRING (10.8%) but substantially recovered by LIGMAP (67.9%). WikiPathways therefore reproduced the complementarity and preferential metabolite contribution to enzyme-containing sets despite different pathway boundaries. Because Reactome and WikiPathways are not fully independent resources, this analysis is best interpreted as a robustness test across alternative curated representations.

**Table 3.** WikiPathways replication using the fixed comparison cohort^a^.

| Network | Class | Number of pathways | One component | At most 2 components | Average Coverage | Largest | Top 2 |
| --- | --- | --- | --- | --- | --- | --- | --- |
| STRING | Non-enzyme | 110 | 20/110 (18.2%) | 54/110 (49.1%) | 52.6% | 56.9% | 79.4% |
| STRING | Enzyme-only | 17 | 0/17 (0.0%) | 11/17 (64.7%) | 10.8% | 45.6% | 87.3% |
| STRING | Mixed | 808 | 35/808 (4.3%) | 94/808 (11.6%) | 59.0% | 48.1% | 62.2% |
| LIGMAP | Non-enzyme | 110 | 14/110 (12.7%) | 46/110 (41.8%) | 45.4% | 55.0% | 77.0% |
| LIGMAP | Enzyme-only | 17 | 8/17 (47.1%) | 16/17 (94.1%) | 67.9% | 78.7% | 98.0% |
| LIGMAP | Mixed | 808 | 96/808 (11.9%) | 272/808 (33.7%) | 77.3% | 78.4% | 87.0% |
| STRING+LIGMAP | Non-enzyme | 110 | 34/110 (30.9%) | 78/110 (70.9%) | 69.9% | 74.9% | 92.1% |
| STRING+LIGMAP | Enzyme-only | 17 | 8/17 (47.1%) | 16/17 (94.1%) | 67.9% | 78.7% | 98.0% |
| STRING+LIGMAP | Mixed | 808 | 337/808 (41.7%) | 557/808 (68.9%) | 88.7% | 88.8% | 95.0% |
| LIGMAP+BioLiP2 | Non-enzyme | 110 | 14/110 (12.7%) | 50/110 (45.5%) | 47.2% | 56.2% | 78.4% |
| LIGMAP+BioLiP2 | Enzyme-only | 17 | 9/17 (52.9%) | 16/17 (94.1%) | 68.6% | 79.4% | 98.0% |
| LIGMAP+BioLiP2 | Mixed | 808 | 122/808 (15.1%) | 323/808 (40.0%) | 79.9% | 80.3% | 88.9% |
| STRING+BioLiP2+LIGMAP | Non-enzyme | 110 | 35/110 (31.8%) | 82/110 (74.5%) | 70.5% | 75.4% | 92.6% |
| STRING+BioLiP2+LIGMAP | Enzyme-only | 17 | 9/17 (52.9%) | 16/17 (94.1%) | 68.6% | 79.4% | 98.0% |
| STRING+BioLiP2+LIGMAP | Mixed | 808 | 365/808 (45.2%) | 595/808 (73.6%) | 89.9% | 89.7% | 95.8% |

<sup>a</sup>Pathways were evaluated using a fixed set of eligible proteins. “One component” and “At most 2 components” report the number and percentage of pathways whose complete eligible protein sets form one or no more than two connected components, respectively. “Average coverage” is the mean fraction of eligible proteins incident to at least one within-pathway edge. “Largest” and “Top 2” are the mean fractions of eligible proteins contained in the largest or two largest connected components. Isolated eligible proteins are retained as components of size one. Network combinations represent unions of the indicated edge sets.

### Incremental prediction, structural-layer ablation, and feasibility

Pathway reconstruction can be inflated by protein degree and dense projection, so we tested incremental information using fivefold group cross-validation that held out complete Reactome pathway identifiers. Positive within-pathway pairs were matched to negatives by replacing one endpoint with a protein outside the pathway but in the same network-degree quintile. In Table 4, AUROC measures the model’s ability to rank within-pathway protein pairs above negative pairs across classification thresholds and AUPRC measures the corresponding precision–recall tradeoff and emphasizes accurate recovery of positive within-pathway pairs. Higher values indicate better predictive performance. Protein degree alone performed at chance (mean AUROC, 0.501 ± 0.002; AUPRC, 0.504 ± 0.005), indicating that degree matching removed the discriminatory contribution of connectivity alone. Adding STRING produced the largest individual improvement (AUROC, 0.588 ± 0.006; AUPRC, 0.651 ± 0.008). LIGMAP alone provided a smaller signal beyond degree (AUROC, 0.525 ± 0.005; AUPRC, 0.528 ± 0.006). Combining LIGMAP with STRING increased performance to an AUROC of 0.600 ± 0.007 and an AUPRC of 0.660 ± 0.007; the paired fold-level AUPRC improvement over degree plus STRING was 0.0094 (95% CI, 0.0021–0.0168; paired t(4) = 3.56, two-sided P = 0.0236). Adding BioLiP2 contacts further increased AUPRC to 0.663 ± 0.008, an additional paired improvement of 0.0029 (95% CI, 0.0014–0.0043; paired t(4) = 5.47, two-sided P = 0.0054). Both planned incremental comparisons remained significant after Holm correction (adjusted P = 0.0236 and 0.0109, respectively). Because the five training sets overlap, these P values quantify consistency across held-out folds rather than independent experimental replication. STRING supplied most of the predictive signal, while LIGMAP made a reproducible but modest complementary contribution.

**Table 4.** Pathway-held-out, degree-matched predictive performance.

| Model | Mean AUROC | SD | Mean AUPRC | SD |
| --- | --- | --- | --- | --- |
| Degree only | 0.501 | 0.002 | 0.504 | 0.005 |
| Degree + STRING | 0.588 | 0.006 | 0.651 | 0.008 |
| Degree + LIGMAP | 0.525 | 0.005 | 0.528 | 0.006 |
| Degree + STRING + LIGMAP | 0.600 | 0.007 | 0.660 | 0.007 |
| Degree + STRING + LIGMAP + BioLiP2 | 0.601 | 0.007 | 0.663 | 0.008 |

Layer ablation showed that the signal was distributed rather than confined to one class of protein–metabolite interaction. Under the same feature-degree range of 2–100, standalone-monomer, dimer-interface, and dimer-noninterface groups contributed 555,649, 82,147, and 548,970 projected edges, respectively. Repulsive COLIG states supplied the largest coverage increment beyond STRING, particularly for enzyme-only pathways, whereas the dimer-interface layer was smaller but nonzero. This ordering is mechanistically plausible but not causal evidence: attractive and repulsive states are predicted occupancy relationships, and the edge counts reflect both biology and feature-degree distributions (Supplementary Table S4). Tissue-expression filters retained substantial pathway organization, and requiring a shared UniProt subcellular-location annotation yielded mean union coverage of 80.8%, 85.1%, and 87.2% for strict non-enzyme, enzyme-only, and mixed pathways. These filters improve biological feasibility but do not establish simultaneous expression, colocalization, concentration, or occupancy in a single cell.

### Combined evidence for metabolite-enabled pathway organization

Figure 2 integrates three complementary levels of evidence for metabolite-enabled protein organization. Panel A shows that LIGMAP-mediated connections reconstruct conventional pathways and complement STRING across Reactome and WikiPathways. Panel B shows that the incremental contribution persists after controlling for protein degree and holding out complete pathway identifiers. Panel C identifies the structural and COLIG prediction layers that supply additional coverage. The recurrence of this signal across pathway resources, pathway classes, and structural layers argues that metabolite-mediated organization is not confined to a small set of canonical metabolic pathways. Together, these analyses demonstrate convergent pathway reconstruction, predictive validation, and mechanistic decomposition while preserving the distinction between network association and causal regulation.

**Figure 2.**
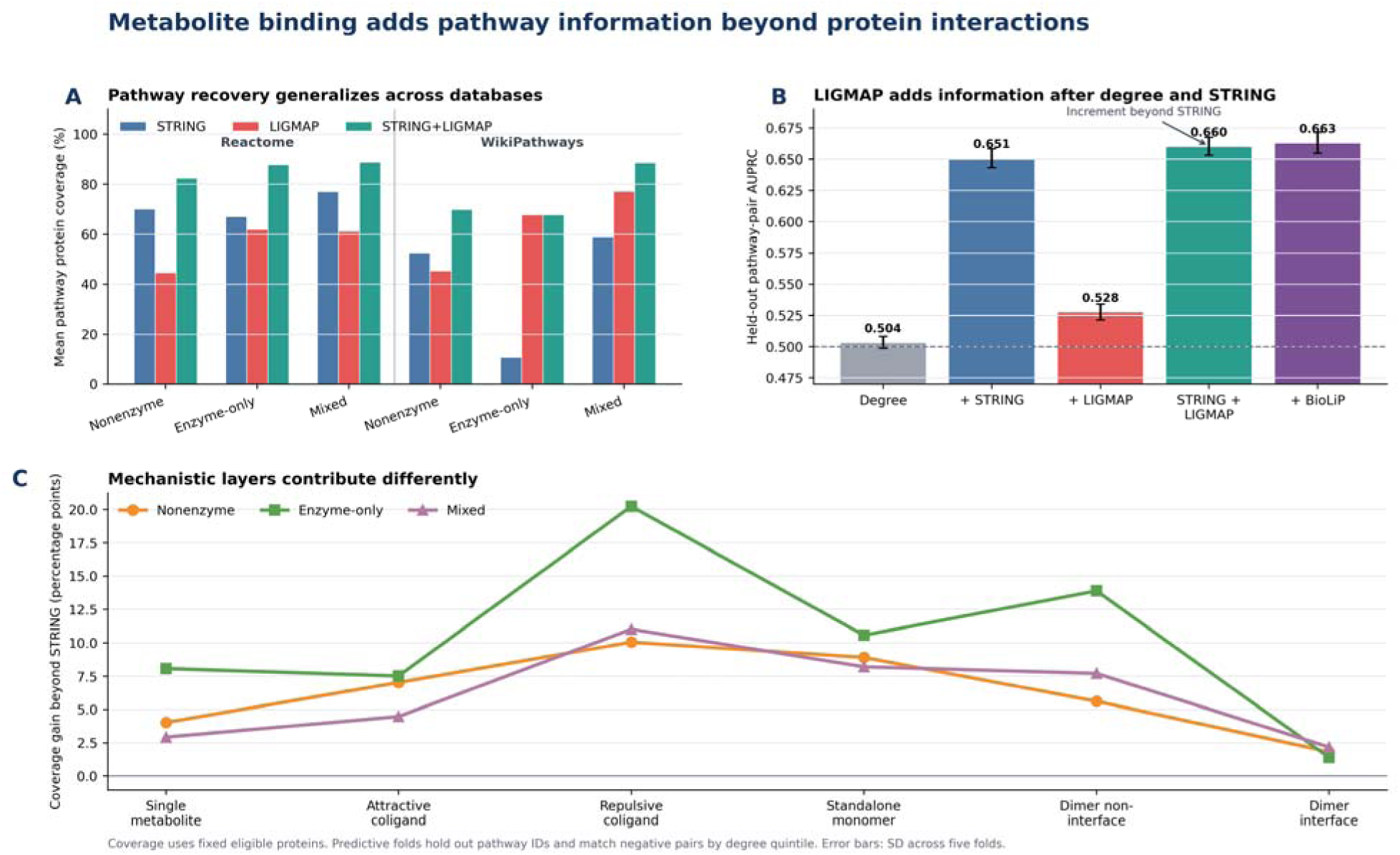
Convergent evidence for metabolite-enabled pathway and global connectivity. (A) Reactome and WikiPathways coverage by STRING, LIGMAP, and their union. (B) Pathway-held-out degree-matched prediction. (C) Layer-specific coverage gains beyond STRING.

### Ancient and non-ancient metabolites make distinct network contributions

Features involving the 34 analyzed ancient metabolites, non-ancient-only metabolites, and the complete retained network were all globally connected or nearly so (Supplementary Table S2A). Their roles nevertheless differed. Ancient-only metabolites yielded 267 projected edges per feature and reached 1,509 proteins; non-ancient-only metabolites yielded 47.6 edges per feature but reached 3,586 proteins. Thus, analyzed ancient metabolites are efficient dense connectors, whereas the larger non-ancient repertoire distributes connectivity more broadly. Removing ancient metabolites reduced pathway coverage but retained substantial organization (Supplementary Table S2B): non-ancient-only mean coverage was 39.9% for strict non-enzyme, 50.1% for enzyme-only, and 57.2% for mixed pathways. Ancient-only values were 13.9%, 19.7%, and 20.7%. Neither partition should be treated as a negative control for the other; both are biologically plausible realizations with different density and breadth.

Table 5 summarizes the global topology generated by ancient and non-ancient retained features. The ancient-only partition contains 504 features connecting 1,509 nodes through 134,801 edges. Although it forms two components, 99.9% of its nodes belong to the largest component, indicating near-complete global connectivity. The non-ancient-only partition is larger, with 15,521 features, 3,586 nodes, and 739,320 edges in a single component. Ancient-involving features generate 552,990 edges among 2,976 nodes and likewise form one component. The complete union contains 22,530 features connecting 3,786 nodes through 951,715 edges, with all nodes in one component. Ancient-only features generate substantially more edges per feature (267.5) than non-ancient-only features (47.6), ancient-involving features (78.9), or the complete union (42.2), showing that ancient-metabolite features are individually associated with especially broad connectivity.

**Table 5.** Global topology of ancient and non-ancient retained features^a^.

| Partition | Features | Nodes | Edges | Components | Largest component | Edges/feature |
| --- | --- | --- | --- | --- | --- | --- |
| Ancient only | 504 | 1509 | 134801 | 2 | 99.9% | 267.5 |
| Non-ancient only | 15521 | 3586 | 739320 | 1 | 100.0% | 47.6 |
| Ancient involving | 7009 | 2976 | 552990 | 1 | 100.0% | 78.9 |
| Union of metabolites | 22530 | 3786 | 951715 | 1 | 100.0% | 42.2 |

<sup>a</sup> Features are retained single-metabolite or exact attractive/repulsive COLIG assignments incident to 2–100 reviewed human proteins. Nodes are proteins assigned to at least one retained feature, and edges are unique undirected protein pairs sharing at least one feature. Components are disconnected groups in the projected protein network, and largest component is the percentage of nodes in the largest such group. Edges/feature is the total number of unique projected edges divided by the number of retained features; because multiple features may support the same edge, it should be interpreted as a network-level edge-to-feature ratio rather than a per-feature mean. Ancient-only features contain exclusively ancient metabolite codes, non-ancient-only features contain none, and ancient-involving features contain at least one ancient code.

Table 6 compares pathway reconstruction by ancient-involving, ancient-only, non-ancient-only, and complete-union feature partitions across enzyme-only, mixed, and non-enzyme pathways. Ancient-only features provide partial reconstruction, with relatively low protein coverage—19.7% for enzyme-only, 20.7% for mixed, and 13.9% for non-enzyme pathways—and few pathways recovered as a single component. The broader ancient-involving partition performs substantially better, reconstructing 27 of 122 enzyme-only, 69 of 1,315 mixed, and 16 of 104 non-enzyme pathways as one component, with 81, 314, and 62 pathways, respectively, reconstructed in at most two components. Non-ancient-only features generally provide greater coverage and connectivity than ancient-only features, particularly for mixed pathways. The complete union of both ancient and non-ancient metabolites produce the strongest overall reconstruction, reaching protein coverages of 54.0%, 63.1%, and 45.1% and reconstructing 41 enzyme-only, 209 mixed, and 29 non-enzyme pathways as single components. It reconstructs 96, 553, and 72 pathways, respectively, in at most two components. These results show that ancient features alone recover a meaningful but limited portion of pathway organization, while ancient and non-ancient features together provide the most comprehensive pathway connectivity.

**Table 6.**
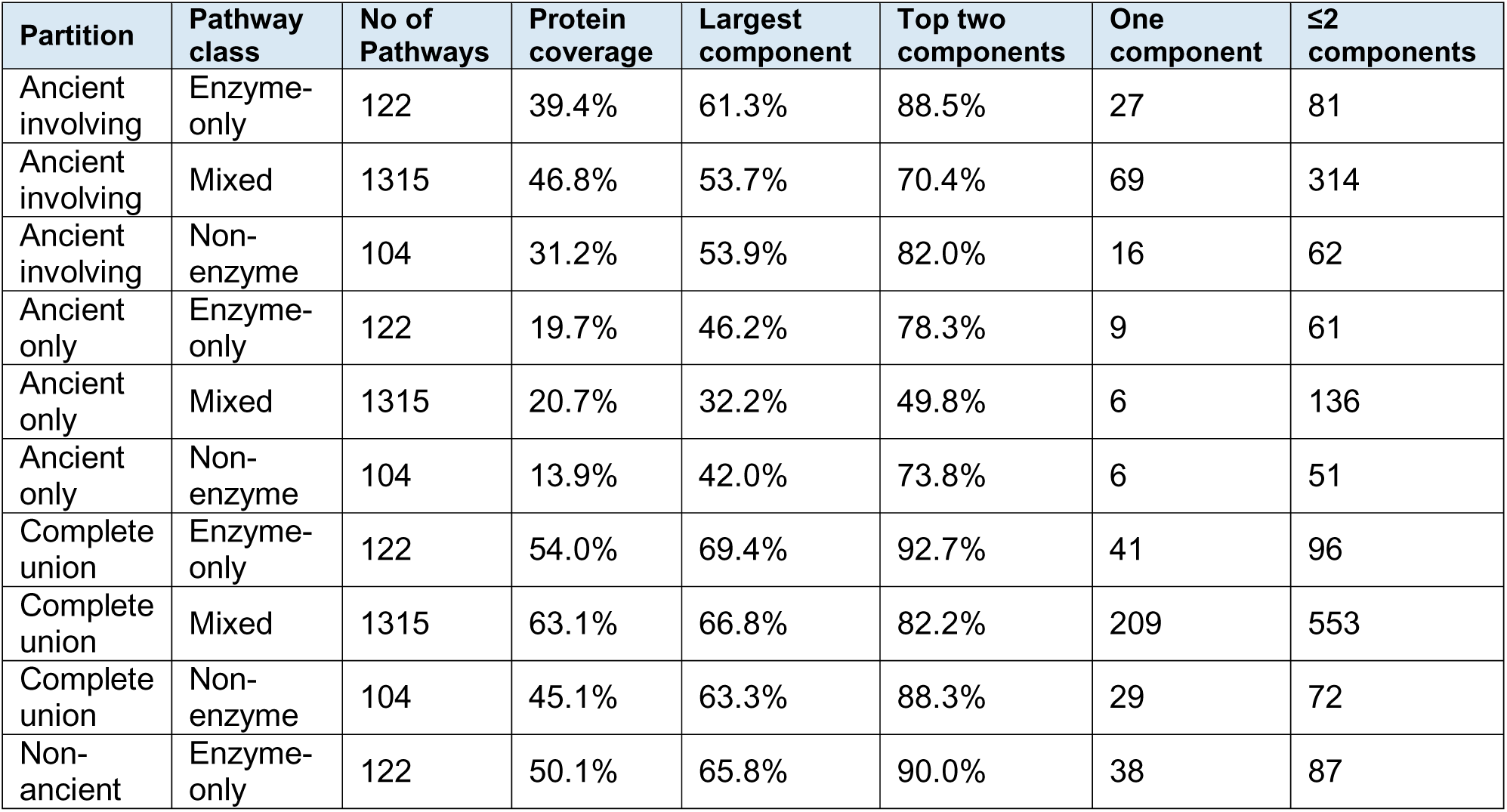
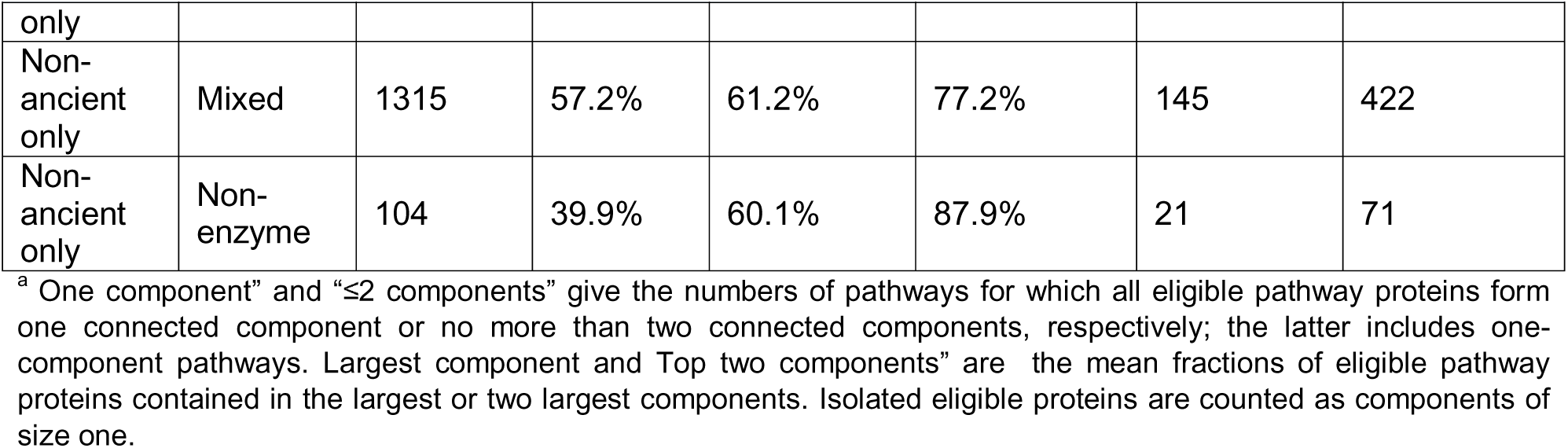
Ancient and non-ancient pathway reconstruction by pathway class^a^.

### Global connectivity is general; specificity is a perturbation of the dense background

At the broader 5,426-protein denominator, cap-100 LIGMAP recovered 698 of 1,691 strict non-enzyme reaction-skeleton edges (41.3%). Protein-label permutation gave a 6.36-fold enrichment (P=0.0001), while the more stringent exact bipartite degree-preserving null retained every protein and feature degree and recovered 640.1 edges on average (observed/null=1.09, P=0.005; Supplementary Table S2C). The observed cap-100 graph contained seven components and a 3,786-protein largest component (99.84% of active proteins); randomized graphs had a mean largest component of 3,782.9 proteins and were likewise nearly global. Thus, the null reproduces the general connectivity-generating property but not all biological placement: the significant 698-versus-640.1 excess is the metabolite-specific component detectable after topology is held fixed. Uncapped LIGMAP recovered 1,668 edges (98.6%) but was indistinguishable from the degree-preserving null (observed/null=0.994), showing that saturation erases this identity-selective signal. Global reach is therefore a general consequence of the incidence-degree architecture, whereas metabolite identity and placement act as a smaller perturbation concentrated among lower-degree features (9, 10, 25).

Table 7 uses a degree-preserving null model to distinguish general network connectedness from pathway-specific metabolite placement. Under the cap-100 condition, the observed network recovered 698 pathway edges compared with a null mean of 640.1, a significant 1.090-fold enrichment (P=0.005). The observed giant connected component (GCC) contained 3,786 proteins, or 99.8% of the active network, closely matching the null mean of 3,782.9. In the uncapped network, 1,668 pathway edges were recovered compared with 1,678.5 under the null (observed/null=0.994; P=0.995), while essentially all 5,426 proteins belonged to the GCC in both observed and randomized networks. Thus, global connectivity is largely explained by network degree structure, whereas restricting the analysis to features assigned to no more than 100 proteins reveals a modest but significant excess of pathway-edge recovery attributable to metabolite placement.

**Table 7.**
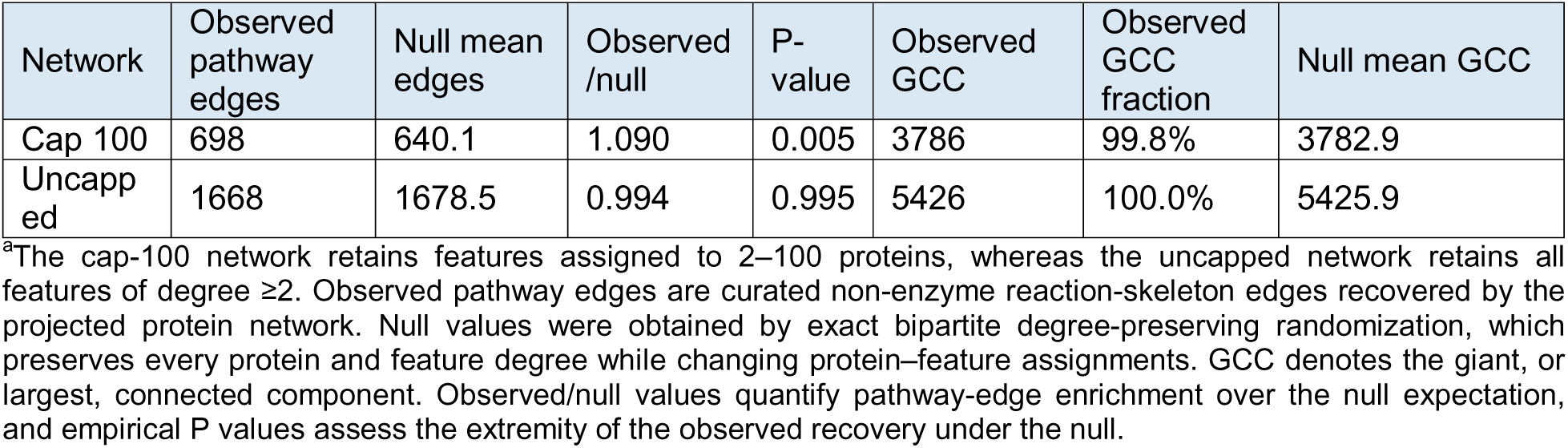
Degree-preserving null separates general connectivity from metabolite-specific placement.

### Sensitivity analyses define where the signal is strongest

The upper feature-degree cap is an explicit selectivity choice, not a biologically known threshold. Raising it monotonically increased descriptive coverage and ultimately approached saturation; the degree-preserving null showed why uncapped recovery is not compelling by itself. Conclusions about STRING-LIGMAP complementarity persisted across caps of 50-250 and across pathway-size strata (Supplementary Table S5). Two independent WikiPathways identifier mappings gave nearly identical union coverage, including among pathways with less than 50% membership overlap with any Reactome pathway, reducing concern that the replication is only a Reactome remapping. However, LIGMAP recovery of within-pathway reference pairs was strongly degree-dependent: 0.5%, 1.5%, 6.1%, and 35.1% across ascending endpoint-degree quartiles. Hub concentration is therefore a substantive limitation even though the degree-matched held-out model retains a small incremental signal.

Experimental BioLiP2 contacts provide a second calibration. Across 2,314 exact human protein-metabolite contacts eligible for comparison, the raw single-metabolite LIGMAP incidence recovered 849 (36.7%). The primary cap-100 projection retained only 28 (1.2%), because the cap excludes broadly assigned features rather than optimizing contact recall. Standalone-monomer and non-interface-in-dimer layers recovered 24.2% and 22.9% of raw contacts; the interface layer recovered 2.8%. This contrast prevents conflating two goals: raw LIGMAP is the relevant object for binding-contact recall, whereas cap-100 is a topology-selective network used to limit saturation. No defensible experimental true-negative ligand-binding panel was available locally, so specificity and precision are not estimated by this audit.

### Experimental systems validate the mechanism independently of LIGMAP

Concept validation and prediction validation provide complementary evidence while also identifying areas for computational improvement. Glucose/FBP perturbation changes aldolase-dependent lysosomal complex assembly and AMPK/ACC phosphorylation (26); methionine/SAM changes SAMTOR–GATOR1–KICSTOR association and mTORC1 output (27); and purine nucleotides glue PPAT to NUDT5, altering purine flux and proliferation (28). Inositol tetraphosphate stabilizes HDAC–corepressor assemblies (29, 30). FH loss/fumarate and heme-responsive systems extend these mechanisms across transcriptional, RNA-processing, redox, and respiratory outputs (31–37). These observations establish localized metabolite-controlled protein organization whether or not LIGMAP recovers the corresponding system. Their mechanistic diversity provides independent support for interpreting the global network as a shared organizational capacity rather than the extrapolation of one exceptional regulatory mechanism.

The LIGMAP benchmark is somewhat more limited. Of seven stringent direct mechanisms, exact metabolite assignment plus union connectivity completely recovered fumarate-KEAP1-NRF2 and heme-REV-ERB-NCOR, partially recovered PPAT-NUDT5 through NUDT5, and missed four. It recovered all three broader serial axes tested: DHT-SHBG-AR, MTA-MTAP-PRMT5-MAT2A, and CD73-adenosine-A2AR. STRING alone connected four direct-mechanism groups but none of the serial axes. Neither static network predicts activation, inhibition, stabilization, disruption, concentration dependence, or compartment. A LIGMAP miss is therefore a failure of coverage or prediction, not evidence against an experimentally established mechanism; conversely, a postdiction does not prove a simultaneous cell-wide Entabolon.

### Highlighted mechanistic postdiction: MTAP-MTA-PRMT5 couples metabolism, RNA processing, and antitumor immunity

LIGMAP assigns the metabolite MTA to both MTAP and PRMT5, while COLIG places PRMT5 in the repulsive MTA|SAM set. This is more specific than pathway concordance: it identifies the perturbed metabolite, the target protein, the physiological competing cofactor, and the predicted incompatibility of their occupancy. Independent biochemical and structural studies established that MTAP loss elevates MTA and that MTA competes with SAM in the PRMT5 cofactor pocket, reducing PRMT5 methyltransferase activity (38–40). Genetic screens showed selective dependence of MTAP-deficient cells on PRMT5 and WDR77 and extended the vulnerable axis to MAT2A and the PRMT5-associated protein RIOK1 (38–40).

This response propagates beyond a single enzyme. In T cells, MTA exposure caused 839 significant proteomic changes; at least 75% of MTA- and selective-PRMT5-inhibitor-responsive transcripts overlapped, with reported directional correlation r2=0.94. The affected programs included symmetric arginine methylation, RNA splicing, cytokine and T-cell-receptor signaling, proliferation, and apoptosis. Enzymatic MTA depletion restored T-cell function and improved checkpoint-immunotherapy responses in MTAP-deficient tumor models (41). This causal, reversible, cross-pathway and multicellular response is consistent with a biologically expressed Entabolon-like state.

This postdiction is kept separate from the capped reconstruction statistics. The single-MTA feature has degree 274, and REP:MTA|SAM has degree 105, narrowly exceeding the prespecified cap of 100. It therefore validates information present in the raw mechanistic predictions but was not used to improve capped pathway recovery. Context also matters: some primary MTAP-deleted glioblastomas fail to accumulate MTA because MTAP-positive stroma clears it (42). That limitation supports a conditional architecture governed by abundance, transport, and neighboring cells rather than a constitutively active global network.

Table 8 compares LIGMAP and STRING methods against ten experimentally established metabolite-dependent regulatory systems. LIGMAP achieved complete target recall and recovered the required metabolite-binding prerequisite in five systems: KEAP1–fumarate– NRF2, REV-ERB–heme–NCOR, SHBG–DHT–AR, MTA–MTAP–PRMT5–MAT2A, and CD73–adenosine–A2AR. The PPAT–NUDT5 purine-glue system showed partial target recall of 50%, whereas the HDAC3–NCOR2, ALDOA–FBP–AMPK, SAMTOR–SAM–GATOR1, and DGCR8–heme systems were not recovered by LIGMAP. STRING alone placed four systems in one component, five in two components, and the MTA system in three components. Overall, the benchmark shows that the network recovers several experimentally supported Entabolon-like mechanisms, including all three serial systems, but misses or only partially captures several direct metabolite-dependent assemblies. These results provide targeted experimental support for the framework while also demonstrating that the current predictions do not comprehensively recover all known mechanisms.

**Table 8.**
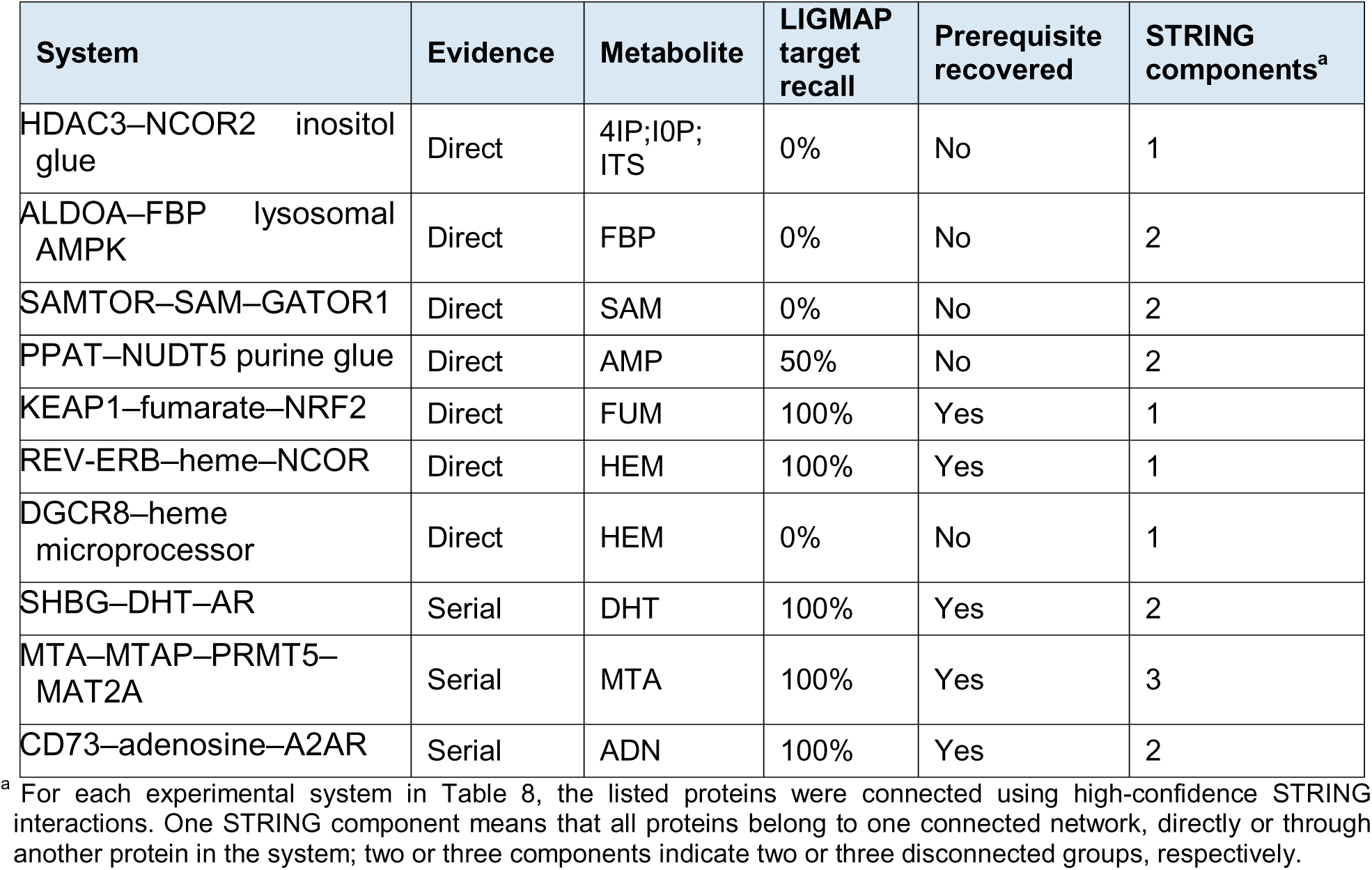
Experimental mechanism benchmark.

| System | Evidence | Metabolite | LIGMAP target recall | Prerequisite recovered | STRING components<sup>a</sup> |
| --- | --- | --- | --- | --- | --- |
| HDAC3–NCOR2 inositol glue | Direct | 4IP;IOP; ITS | 0% | No | 1 |
| ALDOA–FBP lysosomal AMPK | Direct | FBP | 0% | No | 2 |
| SAMTOR–SAM–GATOR1 | Direct | SAM | 0% | No | 2 |
| PPAT–NUDT5 purine glue | Direct | AMP | 50% | No | 2 |
| KEAP1–fumarate–NRF2 | Direct | FUM | 100% | Yes | 1 |
| REV–ERB–heme–NCOR | Direct | HEM | 100% | Yes | 1 |
| DGCR8–heme microprocessor | Direct | HEM | 0% | No | 1 |
| SHBG–DHT–AR | Serial | DHT | 100% | Yes | 2 |
| MTA–MTAP–PRMT5–MAT2A | Serial | MTA | 100% | Yes | 3 |
| CD73–adenosine–A2AR | Serial | ADN | 100% | Yes | 2 |

<sup>a</sup> For each experimental system in Table 8, the listed proteins were connected using high-confidence STRING interactions. One STRING component means that all proteins belong to one connected network, directly or through another protein in the system; two or three components indicate two or three disconnected groups, respectively.

## Discussion

By mapping metabolites to putative or experimentally established binding sites on protein monomers and dimers, a limited metabolite repertoire reconstructed substantial pathway structure and complemented STRING, particularly when enzymes and non-enzymes co-occurred. Largest- and two-component results showed that requiring one perfectly connected component can obscure extensive reconstruction: for strict non-enzyme Reactome pathways, STRING+LIGMAP placed an average of 86.3% of eligible proteins in the largest component and 95.1% in the two largest components. Replication in WikiPathways showed that this conclusion was not peculiar to one set of curated boundaries. Absolute coverage differed between resources because their membership, scale, granularity, and curation models differ; the reproducible observation was the gain from combining metabolite and protein–protein interaction edges.

The deeper implication is that metabolites may generate a global latent coupling architecture. Highly connected ancient metabolites efficiently create dense reach, while non-ancient metabolites independently generate a globally connected network with broader protein coverage (Tables 5 and 6; Supplementary Table S2A, B). Ancient status therefore changes the density and distribution of connectivity, not the qualitative capacity to form a global network. The degree-preserving null sharpens this interpretation (Table 7; Supplementary Table S2C): nearly identical giant components show that global connectivity is an inherent connectivity-generating property of the incidence architecture, whereas the significant cap-100 excess of pathway-aligned edges isolates a smaller metabolite-specific placement component. The null is not a competing biological model; it separates what any network with these degrees can accomplish from the additional alignment contributed by the observed metabolite assignments. Thus, the global capacity is not dependent on one metabolite, pathway, or evolutionary class. Moreover, the null reflects the inherent background (like a circuit of equal resistance) of the putative metabolite connectivity; it gives many of the qualitative features but not the actual quantitative output. Biology could exploit this general connected architecture, while functional selectivity enters through metabolite identity, spatial and temporal availability, concentration, affinity, occupancy, competition, and protein state.

This Entabolon model extends but does not replace the classical metabolon. Metabolons are physical enzyme assemblies associated with substrate channeling (1–6). Entabolons need not be stable modules: a metabolite may stabilize a complex, disrupt a protein–protein interaction, alter a monomer–multimer equilibrium, or couple proteins serially across compartments and pathways (18, 20). Experimentally established examples show that each mechanism exists locally. Demonstrating a cell-wide Entabolon will require a coordinated metabolite perturbation followed in the same cells and time course by quantitative interactome measurements and matched downstream multiomic or functional readouts.

The strongest conclusion is therefore bounded but consequential. Metabolite-binding networks have sufficient reach to reconstruct conventional pathway organization, add information to protein–protein interactions, and provide a plausible physical basis for collective cross-pathway responses. The current data support selected experimentally realized Entabolon-like neighborhoods and a global architectural hypothesis; they do not establish physiological occupancy of every predicted site or simultaneous activation of the complete projected network. Further computational refinement and prospective experimental validation are therefore required.

To our knowledge, this is the first proteome-scale analysis to show that metabolite-binding relationships generate a nearly connected human protein network spanning conventional pathways, protein functional classes, and evolutionary ages. This architecture suggests that metabolites constitute a pervasive coordination layer linking proteins that need not interact directly. Metabolite-mediated protein coordination is therefore not restricted to isolated pathways or exceptional metabolites; rather, the capacity for such coordination emerges as a global organizational property of the protein–metabolite interaction system. The present work establishes the network architecture through which this coordination could occur and defines the prospective perturbation experiments needed to test its dynamic realization in cells.

## Methods

### Proteins and metabolite library

Supplementary Table S1 lists the 308 human metabolite codes considered. The primary network retained metabolite or COLIG features assigned to 2–100 proteins; an uncapped sensitivity analysis retained all features assigned to at least two proteins. STRING used the physical subnetwork at combined score ≥700 (7), and BioLiP2 was restricted to reviewed human proteins and supplied human–metabolite codes (23). The fixed comparison cohort comprised 3,938 proteins represented by at least one retained LIGMAP or BioLiP2 feature. Reactome pathways (8) were classified using UniProt EC annotations (43) as strict non-enzyme, enzyme-only, or mixed. Protein coverage was the fraction of fixed eligible pathway proteins incident to at least one within-pathway edge; therefore, A–B and B–C cover all three proteins. Largest- and top-two-component fractions used the same denominator.

### Protein and pathway annotation

Reviewed human accessions and EC annotations were obtained from UniProt (43). PDB chains were mapped through SIFTS (44), and PDB/PDBe structures supplied dimer assemblies (45, 46). Reactome human memberships and reaction annotations defined the primary pathway proteins and reaction-skeleton reference edges (8). The July 10, 2026 human WikiPathways GMT release supplied an alternative community-curated membership benchmark (24). WikiPathways Entrez Gene identifiers were mapped through STRING aliases to reviewed human UniProt accessions; 23,461 of 40,228 unique within-pathway membership records had at least one reviewed mapping. ChEBI identifiers supported metabolite and reactant/product mapping (47).

### LIGMAP and COLIG construction

CAVITATOR identified candidate pockets, and APoc aligned target pockets to ligand-bound structural templates (18–21). LIGMAP transferred a metabolite when the APoc pocket-alignment P value was ≤0.003, at least six aligned pocket residues were identical, and at least four ligand templates supported the assignment (18). At this operating point, the reported benchmark precision and recall were approximately 0.66 and 0.142, respectively (18). COLIG states represented two metabolites predicted to occupy the same pocket. Attractive pairs satisfied nonclashing-pose and interligand-contact criteria; repulsive pairs represented mutually incompatible occupancy or unfavorable interligand interactions (22). LIGMAP generated single-metabolite, attractive-pair, and repulsive-pair assignments for standalone monomers, protein-interface pockets, and non-interface sites in dimers. Protein structures were obtained from the PDB (46). Unless explicitly identified as BioLiP2 evidence, metabolite-binding assignments were LIGMAP predictions; BioLiP2 supplied experimentally resolved protein–metabolite contacts (23).

### Network projection and method comparison

Metabolite-mediated protein networks were constructed by projecting the protein– metabolite incidence map into a protein–protein network. Each single metabolite or exact attractive or repulsive COLIG state was treated as a separate feature. For every retained feature, all proteins assigned that feature were connected to one another. The neighbors of a given protein were then defined as the union of all other proteins that shared at least one retained feature with it. Thus, if the same two proteins shared several metabolites or COLIG states, their connection was counted as one unweighted protein–protein edge, while the number and identities of the shared features were retained as supporting information. The feature-degree cap was applied to metabolites and COLIG states rather than to individual proteins or their final network degrees. In the primary cap-100 analysis, a feature was retained only when it was assigned to between 2 and 100 proteins. If a metabolite was assigned to 200 proteins, the entire metabolite feature was excluded; no subset of 100 proteins was selected. A protein could nevertheless have more than 100 neighbors if it was connected to different proteins through multiple retained features. Alternative feature-degree thresholds were examined in sensitivity analyses to determine whether the conclusions depended on the selected cap.

STRING v12 physical links with combined score ≥700 provided the protein–protein interaction baseline (7). BioLiP2 contacts were restricted to reviewed human proteins and supplied human–metabolite codes; DNA, RNA, and peptide ligands were excluded (23). Five models were compared on the same 3,938 eligible proteins: STRING, LIGMAP, STRING+LIGMAP, LIGMAP+BioLiP2, and STRING+BioLiP2+LIGMAP.

### Incremental prediction, ablation, and biological-feasibility filters

Incremental prediction used fivefold GroupKFold cross-validation (48), with Reactome pathway identifiers defining held-out groups. Positive pairs comprised eligible within-pathway pairs, limited to 2,000 per pathway; each negative replaced one endpoint with a protein outside the pathway from the same total-network-degree quintile. Logistic regression used log-transformed endpoint degrees and their product, followed by binary STRING, LIGMAP, and BioLiP2 edge indicators. Standardization was fitted within each training fold. AUROC and area under the precision–recall curve were macro-averaged across folds. Planned incremental model comparisons used two-sided paired t-tests on the five matched fold scores; 95% confidence intervals were calculated for paired differences, and the two incremental AUPRC comparisons were Holm-adjusted. Because training sets overlap across folds, these tests assess fold-level consistency rather than independent replication. Layer ablations separately united the three standalone-monomer, dimer-interface, or dimer-noninterface archives and the single-metabolite, attractive, or repulsive feature types, always applying the same feature-degree cap. Tissue filters required Human Protein Atlas consensus expression in the named tissue (49); the localization filter required at least one shared UniProt subcellular-location term (43). These were permissive feasibility screens rather than cell-type-resolved occupancy models.

### Coverage and component statistics

Protein coverage was the fraction of fixed eligible target proteins incident to at least one within-target edge. Connected components included isolated eligible proteins. Largest-component coverage and top-two-component coverage divided the number of proteins in the largest or two largest induced components by the number of fixed eligible target proteins. Mixed pathways were evaluated as complete pathways, non-enzyme cores, cores plus the single eligible pathway enzyme maximizing largest-component coverage, and cores plus all eligible pathway enzymes. Full per-pathway values are supplied as Supplementary Data 1.

### Ancient-metabolite partitions and topology controls

34 ancient metabolites were present in the analyzed 308-metabolite incidence table. The same cap of 100 was applied. Protein-label QAP permutations tested pathway alignment while holding the projected graph fixed. Exact bipartite double-edge swaps preserved every protein degree and every feature degree and separately tested placement (50). Global connectedness and pathway-edge recovery were reported independently.

### Experimental benchmark

Primary studies defined metabolite codes, direct targets, context partners, and measured state changes before network scoring (13, 14, 26–42, 45, 47, 51, 52). Complete recovery required exact LIGMAP assignment of every direct metabolite target and one-component connectivity of the specified group in the LIGMAP+STRING union. Direction was not generally inferred from projected edges. The MTA–SAM case was audited separately because the repulsive COLIG state explicitly predicts mutually incompatible occupancy. Fumarate chemoproteomic overlaps were descriptive because the retained local audit lacks the complete assayed background (32).

### Operational glossary and interpretation boundaries

| Term | Operational definition |
| --- | --- |
| Raw incidence | All protein-feature assignments from the complete nine-archive, 308-code August 22 input before feature-degree filtering or clique projection. |
| Feature | One metabolite code or one exact unordered attractive/repulsive COLIG pair in a specified structural layer. |
| Feature degree | Number of distinct reviewed human proteins assigned to a feature within the stated cohort. |
| Capped / cap 100 | Primary network retaining features with degree 2-100; degree-1 features cannot generate a protein-protein edge and features above 100 are excluded. |
| Uncapped | Sensitivity projection retaining every feature with degree at least 2, with no upper degree limit; it is derived from raw incidence but is not synonymous with the raw table. |
| Projected edge | Undirected protein pair generated because both proteins share a retained metabolite or exact COLIG feature; each feature is projected as a clique. |
| Fixed comparison cohort | The same 3,938 reviewed human proteins represented by at least one retained LIGMAP or BioLiP2 feature, used as the denominator for method comparisons. |
| Eligible pathway protein | A protein that belongs to the evaluated pathway target and to the fixed comparison cohort; eligibility does not require an edge in the tested network. |
| Protein coverage | Fraction of eligible pathway proteins incident to at least one within-pathway edge. Thus A-B plus B-C covers A, B, and C (3/3). |
| Largest/top-two component coverage | Fraction of eligible proteins in the largest, or union of the two largest, connected components of the pathway-induced graph; isolated eligible proteins are components of size one. |
| One component/ $\leq 2$ components | A pathway whose entire eligible target forms one connected component, or no more than two connected components, respectively. |
| Strict non-enzyme/ enzyme-only / mixed | Reactome pathway classes containing, respectively, no EC-annotated proteins, only EC-annotated proteins, or both EC-annotated and EC-negative proteins among the evaluated members. |
| Non-enzyme core | The EC-negative eligible proteins of a mixed pathway, evaluated alone or after adding the specified eligible pathway enzymes. |
| Ancient / non-ancient | Ancient uses the prespecified 34 code list; non-ancient denotes all other codes, not an evolutionary-age inference from network topology. |
| Ancient-only / non-ancient-only feature | A feature whose constituent code(s) are all ancient, or contain no ancient code, respectively; mixed ancient-non-ancient pairs belong to ancient-involving but neither exclusive partition. |
| Pathway reconstruction | Connectivity among fixed eligible proteins of an already annotated pathway, not de novo discovery of the pathway. |
| Entabolon | Metabolite-enabled organization of enzymes and/or non-enzymes that may be transient and need not constitute a stable physical complex. |
| Prediction vs concept validation | LIGMAP recovery tests the prediction system; experimentally demonstrated metabolite-controlled protein association supports the Entabolon concept independently of whether LIGMAP recovers it. |

## Acknowledgments

This research was supported in part by a gift from the Parker H. Petit AI-Driven Drug Discovery Initiative and a grant GM-118039 from the National Institute of General Medical Sciences of the National Institutes of Health. The authors declare no competing interests. We thank Jessica Forness and Rachel Grimley for reviewing this manuscript and Bartosz Ilkowski for his computational support.

## Data and code availability

All analysis programs, fixed-cohort summaries, complete per-pathway coverage tables, ancient/non-ancient partitions, experimental benchmark tables, and the reference library are supplied with the manuscript. Database-derived inputs remain subject to their source licenses and versioning.

