## Supplementary_information for "The human metabolite–protein interactome reveals a global layer of cellular coordination"

*Global metabolite-mediated connectivity recovers human pathway organization*

Numbered tables are labeled exactly as cited in the final manuscript. Supplementary Data 1 is supplied as a separate machine-readable Excel workbook.

Supplementary Table S1 — 308-metabolite inventory

Supplementary Tables S2A–S2C — ancient/non-ancient topology, pathway reconstruction and degree-preserving null

Supplementary Tables S4A–S4C — structural-layer, feature-type and biological-feasibility analyses

Supplementary Tables S5A–S5E — cap, pathway-size, mapping, contact-recall and endpoint-degree sensitivities

Supplementary Data 1 — complete per-pathway component and coverage values

**Supplementary Table S1. Human metabolite codes included in the 308-metabolite LIGMAP library.**

| **PDB CCD code** | **Metabolite name** | **Ancient status** | **Single** | **Attractive COLIG** | **Repulsive COLIG** | **Raw proteins** | **Cap-100 proteins** |
| --- | --- | --- | --- | --- | --- | --- | --- |
| 13P | Name unresolved in source mapping | Nonancient | Yes | Yes | Yes | 71 | 71 |
| 2HA | Glycerone | Nonancient | Yes | No | Yes | 2 | 2 |
| 2OP | Name unresolved in source mapping | Nonancient | Yes | Yes | Yes | 28 | 28 |
| 2PG | 2-Phospho-D-glycerate | Nonancient | Yes | No | Yes | 10 | 10 |
| 3HA | 3-Hydroxyanthranilate | Nonancient | Yes | Yes | Yes | 61 | 61 |
| 3PG | Name unresolved in source mapping | Nonancient | Yes | Yes | Yes | 94 | 94 |
| 5AD | Name unresolved in source mapping | Nonancient | Yes | Yes | Yes | 200 | 188 |
| 5GP | Name unresolved in source mapping | Nonancient | Yes | Yes | Yes | 338 | 289 |
| 5MC | Name unresolved in source mapping | Nonancient | Yes | Yes | Yes | 27 | 27 |
| 5OP | Name unresolved in source mapping | Nonancient | Yes | Yes | Yes | 25 | 25 |
| 6PG | 6-Phospho-D-gluconate | Nonancient | Yes | No | Yes | 9 | 9 |
| 7MG | Name unresolved in source mapping | Nonancient | Yes | No | Yes | 13 | 13 |
| 9CR | Name unresolved in source mapping | Nonancient | Yes | Yes | Yes | 134 | 123 |
| A2G | N-Acetyl-D-glucosamine | Nonancient | Yes | Yes | Yes | 362 | 272 |
| ACD | Arachidonate | Nonancient | Yes | Yes | Yes | 365 | 295 |
| ACH | Name unresolved in source mapping | Nonancient | Yes | Yes | Yes | 26 | 26 |
| ACO | acetyl coenzyme *a | Ancient | Yes | Yes | Yes | 495 | 405 |
| ACY | Name unresolved in source mapping | Nonancient | Yes | Yes | Yes | 2374 | 1130 |
| ADE | Adenine | Nonancient | Yes | Yes | Yes | 709 | 543 |
| ADN | Name unresolved in source mapping | Nonancient | Yes | Yes | Yes | 526 | 433 |
| ADP | adenosine-5'-diphosphate | Ancient | Yes | Yes | Yes | 1937 | 1217 |
| AKG | 2-oxoglutaric acid | Ancient | Yes | Yes | Yes | 715 | 534 |
| ALA | alanine | Ancient | Yes | Yes | Yes | 286 | 220 |
| ALE | L-Adrenaline | Nonancient | Yes | Yes | Yes | 6 | 6 |
| ALY | Name unresolved in source mapping | Nonancient | Yes | Yes | Yes | 867 | 544 |
| AMP | adenosine monophosphate | Ancient | Yes | Yes | Yes | 1238 | 805 |
| AND | Dehydroepiandrosterone | Nonancient | Yes | Yes | Yes | 45 | 45 |
| APR | Name unresolved in source mapping | Nonancient | Yes | Yes | Yes | 381 | 318 |
| ARG | L-Arginine | Nonancient | Yes | Yes | Yes | 460 | 340 |
| ASC | ascorbic acid | Ancient | Yes | Yes | Yes | 418 | 329 |
| ASD | Androstenedione | Nonancient | Yes | Yes | Yes | 264 | 246 |
| ASN | L-Asparagine | Nonancient | Yes | No | Yes | 13 | 13 |
| ASO | Name unresolved in source mapping | Nonancient | Yes | Yes | Yes | 36 | 36 |
| ASP | aspartic acid | Ancient | Yes | Yes | Yes | 324 | 244 |
| ATP | adenosine-5'-triphosphate | Ancient | Yes | Yes | Yes | 1622 | 1051 |
| AYA | Name unresolved in source mapping | Nonancient | Yes | Yes | Yes | 31 | 31 |
| B12 | Name unresolved in source mapping | Nonancient | Yes | Yes | Yes | 168 | 156 |
| BDF | D-Glucose | Nonancient | Yes | Yes | Yes | 313 | 182 |
| BGC | Name unresolved in source mapping | Nonancient | Yes | Yes | Yes | 916 | 561 |
| BGP | D-Glucose 6-phosphate | Nonancient | Yes | Yes | Yes | 13 | 13 |
| BLA | Name unresolved in source mapping | Nonancient | Yes | Yes | Yes | 437 | 399 |
| BLV | Name unresolved in source mapping | Nonancient | Yes | Yes | Yes | 9 | 9 |
| BMA | Name unresolved in source mapping | Nonancient | Yes | Yes | Yes | 188 | 145 |
| BTN | biotin | Ancient | Yes | Yes | Yes | 194 | 168 |
| C2F | Name unresolved in source mapping | Nonancient | Yes | Yes | Yes | 11 | 11 |
| CAA | Name unresolved in source mapping | Nonancient | Yes | Yes | Yes | 15 | 15 |
| CDP | Name unresolved in source mapping | Nonancient | Yes | Yes | Yes | 139 | 126 |
| CGU | Name unresolved in source mapping | Nonancient | Yes | Yes | Yes | 96 | 96 |
| CHD | Cholic acid | Nonancient | Yes | Yes | Yes | 205 | 193 |
| CHO | Name unresolved in source mapping | Nonancient | Yes | Yes | Yes | 113 | 108 |
| CHT | Name unresolved in source mapping | Nonancient | Yes | Yes | Yes | 347 | 242 |
| CIR | L-Citrulline | Nonancient | Yes | Yes | Yes | 52 | 52 |
| CIT | citric acid | Ancient | Yes | Yes | Yes | 2426 | 1161 |
| CLR | Cholesterol | Nonancient | Yes | Yes | Yes | 923 | 593 |
| CMO | Name unresolved in source mapping | Nonancient | Yes | Yes | Yes | 1072 | 789 |
| CMP | adenosine-3',5'-cyclic-monophosphate | Ancient | Yes | Yes | Yes | 396 | 358 |
| CNC | Name unresolved in source mapping | Nonancient | Yes | Yes | Yes | 49 | 49 |
| CO2 | Name unresolved in source mapping | Nonancient | Yes | Yes | Yes | 552 | 342 |
| CO3 | Name unresolved in source mapping | Nonancient | Yes | Yes | Yes | 986 | 508 |
| COA | coenzyme a | Ancient | Yes | Yes | Yes | 1033 | 719 |
| COB | Name unresolved in source mapping | Nonancient | Yes | No | Yes | 5 | 5 |
| COO | Name unresolved in source mapping | Nonancient | Yes | Yes | Yes | 26 | 26 |
| CSD | Name unresolved in source mapping | Nonancient | Yes | Yes | Yes | 1447 | 807 |
| CSP | Name unresolved in source mapping | Nonancient | Yes | Yes | Yes | 185 | 142 |
| CSX | Name unresolved in source mapping | Nonancient | Yes | Yes | Yes | 891 | 536 |
| CTN | Name unresolved in source mapping | Nonancient | Yes | Yes | Yes | 47 | 47 |
| CTP | Name unresolved in source mapping | Nonancient | Yes | Yes | Yes | 123 | 112 |
| CU1 | Name unresolved in source mapping | Nonancient | Yes | No | Yes | 364 | 172 |
| CYS | L-Cysteine | Nonancient | Yes | Yes | Yes | 462 | 341 |
| CYT | Cytosine | Nonancient | Yes | No | Yes | 10 | 10 |
| DA2 | NG,NG-Dimethyl-L-arginine | Nonancient | Yes | No | Yes | 4 | 4 |
| DCA | Name unresolved in source mapping | Nonancient | Yes | Yes | Yes | 4 | 4 |
| DCP | Name unresolved in source mapping | Nonancient | Yes | Yes | Yes | 48 | 48 |
| DGP | dGMP | Nonancient | Yes | Yes | Yes | 36 | 36 |
| DGT | dGTP | Nonancient | Yes | Yes | Yes | 80 | 80 |
| DHB | 3,4-Dihydroxybenzoate | Nonancient | Yes | Yes | Yes | 70 | 70 |
| DHF | Name unresolved in source mapping | Nonancient | Yes | Yes | Yes | 117 | 104 |
| DHT | Androsterone | Nonancient | Yes | Yes | Yes | 420 | 351 |
| DUR | Deoxyuridine | Nonancient | Yes | Yes | Yes | 12 | 12 |
| DUT | dUTP | Nonancient | Yes | No | Yes | 7 | 7 |
| DXC | 3-Oxo-5beta-cholanate | Nonancient | Yes | Yes | Yes | 335 | 297 |
| EIC | Linoleate | Nonancient | Yes | Yes | Yes | 123 | 121 |
| EPA | (5Z,8Z,11Z,14Z,17Z)-Icosapentaenoic acid | Nonancient | Yes | Yes | Yes | 82 | 82 |
| EST | Estradiol-17beta | Nonancient | Yes | Yes | Yes | 307 | 279 |
| F3S | Name unresolved in source mapping | Nonancient | Yes | Yes | Yes | 381 | 292 |
| F6P | D-Fructose 6-phosphate | Nonancient | Yes | Yes | Yes | 150 | 143 |
| FAD | flavin-adenine dinucleotide | Ancient | Yes | Yes | Yes | 1427 | 1073 |
| FAR | Squalene | Nonancient | Yes | Yes | Yes | 61 | 61 |
| FBP | 1,6-di-o-phosphono-beta-d-fructofuranose | Ancient | Yes | Yes | Yes | 58 | 58 |
| FDP | beta-D-Fructose 2,6-bisphosphate | Nonancient | Yes | Yes | Yes | 21 | 21 |
| FE2 | Name unresolved in source mapping | Nonancient | Yes | No | Yes | 278 | 149 |
| FEC | Name unresolved in source mapping | Nonancient | Yes | Yes | Yes | 13 | 13 |
| FES | Name unresolved in source mapping | Nonancient | Yes | Yes | Yes | 686 | 426 |
| FFO | Name unresolved in source mapping | Nonancient | Yes | Yes | Yes | 5 | 5 |
| FME | Name unresolved in source mapping | Nonancient | Yes | Yes | Yes | 260 | 172 |
| FMN | flavin mononucleotide | Ancient | Yes | Yes | Yes | 1660 | 1037 |
| FOL | Folate | Nonancient | Yes | Yes | Yes | 328 | 294 |
| FPP | Geranylgeranyl diphosphate | Nonancient | Yes | Yes | Yes | 28 | 28 |
| FRU | D-Fructose | Nonancient | Yes | Yes | Yes | 222 | 192 |
| FUC | Name unresolved in source mapping | Nonancient | Yes | Yes | Yes | 261 | 171 |
| FUL | Name unresolved in source mapping | Nonancient | Yes | Yes | Yes | 150 | 122 |
| FUM | fumaric acid | Ancient | Yes | Yes | Yes | 223 | 185 |
| G1P | D-Glucose 1-phosphate | Nonancient | Yes | No | Yes | 28 | 28 |
| G3H | sn-Glycerol 3-phosphate | Nonancient | Yes | Yes | Yes | 24 | 24 |
| G3P | sn-Glycerol 3-phosphate | Nonancient | Yes | Yes | Yes | 187 | 165 |
| G6P | D-Glucose 6-phosphate | Nonancient | Yes | Yes | Yes | 54 | 54 |
| GAB | Gabaculine | Nonancient | Yes | Yes | Yes | 288 | 252 |
| GAL | Name unresolved in source mapping | Nonancient | Yes | Yes | Yes | 323 | 265 |
| GCO | Name unresolved in source mapping | Nonancient | Yes | Yes | Yes | 27 | 27 |
| GCS | D-Glucosamine | Nonancient | Yes | Yes | Yes | 71 | 71 |
| GCU | Name unresolved in source mapping | Nonancient | Yes | Yes | Yes | 23 | 23 |
| GDP | guanosine-5'-diphosphate | Ancient | Yes | Yes | Yes | 1770 | 1025 |
| GDS | Glutathione disulfide | Nonancient | Yes | Yes | Yes | 35 | 35 |
| GLA | Name unresolved in source mapping | Nonancient | Yes | Yes | Yes | 60 | 60 |
| GLC | D-Glucose | Nonancient | Yes | Yes | Yes | 840 | 543 |
| GLN | L-Glutamine | Nonancient | Yes | Yes | Yes | 138 | 112 |
| GLO | Name unresolved in source mapping | Nonancient | Yes | Yes | Yes | 11 | 11 |
| GLP | D-Glucosamine 6-phosphate | Nonancient | Yes | Yes | Yes | 78 | 78 |
| GLU | glutamic acid | Ancient | Yes | Yes | Yes | 400 | 315 |
| GLV | Name unresolved in source mapping | Nonancient | Yes | Yes | Yes | 98 | 98 |
| GLY | glycine | Ancient | Yes | Yes | Yes | 693 | 458 |
| GMP | guanosine | Ancient | Yes | Yes | Yes | 232 | 198 |
| GPP | Geranyl diphosphate | Nonancient | Yes | Yes | Yes | 145 | 129 |
| GSE | Name unresolved in source mapping | Nonancient | Yes | No | Yes | 17 | 17 |
| GSH | glutathione | Ancient | Yes | Yes | Yes | 1196 | 671 |
| GTP | guanosine-5'-triphosphate | Ancient | Yes | Yes | Yes | 645 | 500 |
| GUA | Glutarate | Nonancient | Yes | Yes | Yes | 41 | 41 |
| GUN | Guanine | Nonancient | Yes | Yes | Yes | 39 | 39 |
| H2U | Name unresolved in source mapping | Nonancient | Yes | No | Yes | 3 | 3 |
| H4B | Name unresolved in source mapping | Nonancient | Yes | Yes | Yes | 148 | 140 |
| HAR | N-(omega)-Hydroxyarginine | Nonancient | Yes | Yes | Yes | 48 | 48 |
| HCD | Cholesterol | Nonancient | Yes | Yes | Yes | 73 | 73 |
| HCY | Cortisol | Nonancient | Yes | Yes | Yes | 70 | 70 |
| HEC | Name unresolved in source mapping | Nonancient | Yes | Yes | Yes | 1690 | 1163 |
| HEM | protoporphyrin ix containing fe | Ancient | Yes | Yes | Yes | 2489 | 1580 |
| HIP | Name unresolved in source mapping | Nonancient | Yes | No | Yes | 35 | 35 |
| HIS | L-Histidine | Nonancient | Yes | Yes | Yes | 444 | 358 |
| HMG | Name unresolved in source mapping | Nonancient | Yes | No | Yes | 10 | 10 |
| HPA | Hypoxanthine | Nonancient | Yes | Yes | Yes | 88 | 88 |
| HSE | L-Homoserine | Nonancient | Yes | Yes | Yes | 104 | 100 |
| HSM | Name unresolved in source mapping | Nonancient | Yes | Yes | Yes | 137 | 134 |
| HSX | Name unresolved in source mapping | Nonancient | Yes | Yes | Yes | 5 | 5 |
| HXA | (4Z,7Z,10Z,13Z,16Z,19Z)-Docosahexaenoic acid | Nonancient | Yes | Yes | Yes | 117 | 107 |
| HY3 | Name unresolved in source mapping | Nonancient | Yes | Yes | Yes | 47 | 47 |
| HYP | Name unresolved in source mapping | Nonancient | Yes | Yes | Yes | 101 | 69 |
| ICT | Citrate | Nonancient | Yes | Yes | Yes | 26 | 26 |
| IHP | Name unresolved in source mapping | Nonancient | Yes | Yes | Yes | 504 | 368 |
| ILE | L-Leucine | Nonancient | Yes | Yes | Yes | 71 | 71 |
| IMP | Name unresolved in source mapping | Nonancient | Yes | Yes | Yes | 356 | 297 |
| INS | muco-Inositol | Nonancient | Yes | Yes | Yes | 126 | 100 |
| IPE | Dimethylallyl diphosphate | Nonancient | Yes | Yes | Yes | 330 | 304 |
| IYR | 3-Iodo-L-tyrosine | Nonancient | Yes | Yes | Yes | 316 | 214 |
| KYN | L-Kynurenine | Nonancient | Yes | Yes | Yes | 14 | 14 |
| LAC | Name unresolved in source mapping | Nonancient | Yes | Yes | Yes | 107 | 89 |
| LBN | Name unresolved in source mapping | Nonancient | Yes | Yes | Yes | 74 | 74 |
| LDP | Catecholamine | Nonancient | Yes | Yes | Yes | 110 | 107 |
| LEU | L-Leucine | Nonancient | Yes | Yes | Yes | 262 | 225 |
| LLP | Name unresolved in source mapping | Nonancient | Yes | Yes | Yes | 501 | 408 |
| LMR | (S)-Malate | Nonancient | Yes | Yes | Yes | 345 | 244 |
| LNL | (9Z)-Octadecenoic acid | Nonancient | Yes | Yes | Yes | 110 | 109 |
| LNR | L-Noradrenaline | Nonancient | Yes | Yes | Yes | 144 | 120 |
| LPA | lipoic acid | Ancient | Yes | Yes | Yes | 14 | 14 |
| LPC | 1-Oleoylglycerophosphocholine | Nonancient | Yes | Yes | Yes | 204 | 163 |
| LPE | 1-O-Hexadecyl-lyso-sn-glycero-3-phosphocholine | Nonancient | Yes | Yes | Yes | 45 | 45 |
| LYS | L-Lysine | Nonancient | Yes | Yes | Yes | 445 | 380 |
| M3L | N6-Methyl-L-lysine | Nonancient | Yes | Yes | Yes | 399 | 266 |
| M6P | D-Glucose 6-phosphate | Nonancient | Yes | No | Yes | 5 | 5 |
| M7G | Name unresolved in source mapping | Nonancient | Yes | Yes | Yes | 179 | 163 |
| MAN | Name unresolved in source mapping | Nonancient | Yes | Yes | Yes | 1243 | 670 |
| MC3 | Name unresolved in source mapping | Nonancient | Yes | Yes | Yes | 338 | 265 |
| MET | L-Methionine | Nonancient | Yes | Yes | Yes | 446 | 368 |
| MGD | Name unresolved in source mapping | Nonancient | Yes | Yes | Yes | 81 | 81 |
| MHO | L-Methionine S-oxide | Nonancient | Yes | Yes | Yes | 549 | 407 |
| MLT | (S)-Malate | Nonancient | Yes | Yes | Yes | 487 | 331 |
| MLY | N6-Methyl-L-lysine | Nonancient | Yes | Yes | Yes | 2232 | 1175 |
| MLZ | N6-Methyl-L-lysine | Nonancient | Yes | Yes | Yes | 1009 | 615 |
| MOO | Name unresolved in source mapping | Nonancient | Yes | No | Yes | 75 | 75 |
| MTA | Name unresolved in source mapping | Nonancient | Yes | Yes | Yes | 274 | 243 |
| MTE | Name unresolved in source mapping | Nonancient | Yes | Yes | Yes | 78 | 78 |
| MYR | Hexadecanal | Nonancient | Yes | Yes | Yes | 1605 | 966 |
| NAD | nicotinamide-adenine-dinucleotide | Ancient | Yes | Yes | Yes | 1020 | 808 |
| NAG | Name unresolved in source mapping | Nonancient | Yes | Yes | Yes | 3956 | 1744 |
| NAI | Name unresolved in source mapping | Nonancient | Yes | Yes | Yes | 397 | 361 |
| NAJ | Name unresolved in source mapping | Nonancient | Yes | Yes | Yes | 43 | 43 |
| NAP | nadp nicotinamide-adenine-dinucleotide phosphate | Ancient | Yes | Yes | Yes | 1248 | 909 |
| NCA | Name unresolved in source mapping | Nonancient | Yes | Yes | Yes | 441 | 370 |
| NCN | Name unresolved in source mapping | Nonancient | Yes | No | Yes | 6 | 6 |
| NDG | Name unresolved in source mapping | Nonancient | Yes | Yes | Yes | 410 | 266 |
| NDP | Name unresolved in source mapping | Nonancient | Yes | Yes | Yes | 880 | 696 |
| NEP | Name unresolved in source mapping | Nonancient | Yes | Yes | Yes | 32 | 32 |
| NGA | N-Acetyl-D-glucosamine | Nonancient | Yes | Yes | Yes | 53 | 53 |
| NGE | N-Glycoloyl-neuraminate | Nonancient | Yes | Yes | Yes | 10 | 10 |
| NGY | Name unresolved in source mapping | Nonancient | Yes | Yes | Yes | 7 | 7 |
| NHF | Name unresolved in source mapping | Nonancient | Yes | Yes | Yes | 3 | 3 |
| NIO | Name unresolved in source mapping | Nonancient | Yes | Yes | Yes | 347 | 296 |
| NIY | Name unresolved in source mapping | Nonancient | Yes | Yes | Yes | 317 | 246 |
| NME | Name unresolved in source mapping | Nonancient | Yes | Yes | Yes | 22 | 22 |
| NMN | Name unresolved in source mapping | Nonancient | Yes | Yes | Yes | 144 | 118 |
| OAA | Oxaloacetate | Nonancient | Yes | Yes | Yes | 210 | 194 |
| OLA | (9Z)-Octadecenoic acid | Nonancient | Yes | Yes | Yes | 1939 | 1126 |
| OLC | Name unresolved in source mapping | Nonancient | Yes | Yes | Yes | 2273 | 1219 |
| OMT | Name unresolved in source mapping | Nonancient | Yes | Yes | Yes | 105 | 85 |
| ORN | L-Ornithine | Nonancient | Yes | Yes | Yes | 136 | 106 |
| OXD | Name unresolved in source mapping | Nonancient | Yes | Yes | Yes | 123 | 112 |
| OXL | Name unresolved in source mapping | Nonancient | Yes | Yes | Yes | 206 | 183 |
| OXY | Name unresolved in source mapping | Nonancient | Yes | Yes | Yes | 906 | 648 |
| PA1 | Name unresolved in source mapping | Nonancient | Yes | Yes | Yes | 3 | 3 |
| PAM | (9Z)-Octadecenoic acid | Nonancient | Yes | Yes | Yes | 454 | 339 |
| PAP | Name unresolved in source mapping | Nonancient | Yes | No | Yes | 5 | 5 |
| PAU | Pantothenate | Nonancient | Yes | Yes | Yes | 32 | 32 |
| PCA | Name unresolved in source mapping | Nonancient | Yes | Yes | Yes | 998 | 468 |
| PCF | Name unresolved in source mapping | Nonancient | Yes | Yes | Yes | 66 | 66 |
| PCG | Name unresolved in source mapping | Nonancient | Yes | Yes | Yes | 123 | 111 |
| PCR | Name unresolved in source mapping | Nonancient | Yes | Yes | Yes | 40 | 40 |
| PCW | 1-Hexadecanoyl-2-(9Z-octadecenoyl)-sn-glycero-3-phosphocholine | Nonancient | Yes | Yes | Yes | 304 | 215 |
| PEA | Name unresolved in source mapping | Nonancient | Yes | Yes | Yes | 115 | 110 |
| PEP | phosphoenolpyruvate | Ancient | Yes | Yes | Yes | 98 | 98 |
| PGA | Name unresolved in source mapping | Nonancient | Yes | Yes | Yes | 117 | 104 |
| PGD | Name unresolved in source mapping | Nonancient | Yes | Yes | Yes | 28 | 28 |
| PGM | Name unresolved in source mapping | Nonancient | Yes | No | Yes | 12 | 12 |
| PGP | Name unresolved in source mapping | Nonancient | Yes | Yes | Yes | 12 | 12 |
| PHE | L-Phenylalanine | Nonancient | Yes | Yes | Yes | 265 | 236 |
| PLC | Name unresolved in source mapping | Nonancient | Yes | Yes | Yes | 87 | 87 |
| PLM | Hexadecanal | Nonancient | Yes | Yes | Yes | 1963 | 1150 |
| PLP | pyridoxal-5'-phosphate | Ancient | Yes | Yes | Yes | 506 | 409 |
| PMP | Pyridoxamine phosphate | Nonancient | Yes | Yes | Yes | 54 | 54 |
| PNS | Pantetheine 4'-phosphate | Nonancient | Yes | Yes | Yes | 186 | 145 |
| PO4 | Name unresolved in source mapping | Nonancient | Yes | Yes | Yes | 4153 | 1668 |
| POV | 1-Hexadecanoyl-2-(9Z-octadecenoyl)-sn-glycero-3-phosphocholine | Nonancient | Yes | Yes | Yes | 79 | 79 |
| PP9 | Name unresolved in source mapping | Nonancient | Yes | Yes | Yes | 69 | 69 |
| PPI | Name unresolved in source mapping | Nonancient | Yes | Yes | Yes | 281 | 191 |
| PPS | Name unresolved in source mapping | Nonancient | Yes | Yes | Yes | 36 | 36 |
| PPY | Phenylpyruvate | Nonancient | Yes | Yes | Yes | 56 | 56 |
| PRO | Name unresolved in source mapping | Nonancient | Yes | Yes | Yes | 416 | 314 |
| PSC | Name unresolved in source mapping | Nonancient | Yes | Yes | Yes | 170 | 139 |
| PTR | Phosphotyrosine | Nonancient | Yes | Yes | Yes | 1201 | 680 |
| PTY | Name unresolved in source mapping | Nonancient | Yes | Yes | Yes | 222 | 189 |
| PX4 | Name unresolved in source mapping | Nonancient | Yes | Yes | Yes | 443 | 316 |
| PXP | Pyridoxine phosphate | Nonancient | Yes | No | Yes | 9 | 9 |
| PYR | pyruvic acid | Ancient | Yes | Yes | Yes | 275 | 205 |
| R5P | D-Arabinose 5-phosphate | Nonancient | Yes | Yes | Yes | 8 | 8 |
| RBF | riboflavin | Ancient | Yes | Yes | Yes | 187 | 138 |
| REA | Retinoate | Nonancient | Yes | Yes | Yes | 156 | 150 |
| RED | Name unresolved in source mapping | Nonancient | Yes | No | Yes | 4 | 4 |
| RET | Rhodopsin | Nonancient | Yes | Yes | Yes | 1597 | 1069 |
| RIB | D-Ribulose | Nonancient | Yes | Yes | Yes | 39 | 39 |
| RIP | 1,4-beta-D-Xylan | Nonancient | Yes | Yes | Yes | 224 | 149 |
| RP5 | Name unresolved in source mapping | Nonancient | Yes | Yes | Yes | 11 | 11 |
| RPI | Name unresolved in source mapping | Nonancient | Yes | Yes | Yes | 66 | 66 |
| RTL | Name unresolved in source mapping | Nonancient | Yes | Yes | Yes | 224 | 200 |
| S1P | Sphingosine 1-phosphate | Nonancient | Yes | Yes | Yes | 52 | 52 |
| S7P | Name unresolved in source mapping | Nonancient | Yes | No | Yes | 2 | 2 |
| SAC | Name unresolved in source mapping | Nonancient | Yes | Yes | Yes | 21 | 21 |
| SAH | s-adenosyl-l-homocysteine | Ancient | Yes | Yes | Yes | 1268 | 851 |
| SAM | s-adenosylmethionine | Ancient | Yes | Yes | Yes | 896 | 687 |
| SAR | Name unresolved in source mapping | Nonancient | Yes | Yes | Yes | 103 | 93 |
| SCA | Name unresolved in source mapping | Nonancient | Yes | Yes | Yes | 46 | 46 |
| SEC | Name unresolved in source mapping | Nonancient | Yes | Yes | Yes | 103 | 95 |
| SEP | O-Phospho-L-serine | Nonancient | Yes | Yes | Yes | 1992 | 829 |
| SER | serine | Ancient | Yes | Yes | Yes | 286 | 221 |
| SF4 | Name unresolved in source mapping | Nonancient | Yes | Yes | Yes | 1694 | 893 |
| SGN | N,6-O-Disulfo-D-glucosamine | Nonancient | Yes | No | Yes | 1 | 0 |
| SIA | Name unresolved in source mapping | Nonancient | Yes | Yes | Yes | 209 | 150 |
| SLB | N-Acetylneuraminate | Nonancient | Yes | Yes | Yes | 129 | 117 |
| SMN | (R)-Mandelate | Nonancient | Yes | Yes | Yes | 34 | 34 |
| SNN | Name unresolved in source mapping | Nonancient | Yes | Yes | Yes | 267 | 145 |
| SO4 | Name unresolved in source mapping | Nonancient | Yes | Yes | Yes | 5285 | 1993 |
| SOR | Mannitol | Nonancient | Yes | Yes | Yes | 129 | 96 |
| SPD | Spermidine | Nonancient | Yes | Yes | Yes | 123 | 95 |
| SPH | Hexadecasphinganine | Nonancient | Yes | Yes | Yes | 51 | 51 |
| SPM | Spermidine | Nonancient | Yes | Yes | Yes | 76 | 76 |
| SRO | Serotonin | Nonancient | Yes | Yes | Yes | 243 | 201 |
| STE | Hexadecanoic acid | Nonancient | Yes | Yes | Yes | 443 | 360 |
| T44 | Thyroxine | Nonancient | Yes | Yes | Yes | 165 | 144 |
| TAU | Name unresolved in source mapping | Nonancient | Yes | Yes | Yes | 27 | 27 |
| TES | Testosterone | Nonancient | Yes | Yes | Yes | 268 | 228 |
| THG | Name unresolved in source mapping | Nonancient | Yes | Yes | Yes | 75 | 75 |
| THM | Thymidine | Nonancient | Yes | Yes | Yes | 137 | 120 |
| THP | Name unresolved in source mapping | Nonancient | Yes | Yes | Yes | 707 | 517 |
| THR | L-Threonine | Nonancient | Yes | Yes | Yes | 53 | 53 |
| TMP | dTMP | Nonancient | Yes | Yes | Yes | 173 | 158 |
| TPO | L-Threonine O-3-phosphate | Nonancient | Yes | Yes | Yes | 1385 | 695 |
| TPP | thiamine diphosphate | Ancient | Yes | Yes | Yes | 58 | 58 |
| TPQ | Name unresolved in source mapping | Nonancient | Yes | No | Yes | 2 | 2 |
| TPS | Name unresolved in source mapping | Nonancient | Yes | Yes | Yes | 11 | 11 |
| TRP | L-Tryptophan | Nonancient | Yes | Yes | Yes | 522 | 379 |
| TTP | dTTP | Nonancient | Yes | Yes | Yes | 122 | 99 |
| TXA | Name unresolved in source mapping | Nonancient | Yes | Yes | Yes | 6 | 6 |
| TYI | 3,5-Diiodo-L-tyrosine | Nonancient | Yes | Yes | Yes | 211 | 107 |
| TYR | L-Tyrosine | Nonancient | Yes | Yes | Yes | 102 | 94 |
| U10 | Ubiquinone-9 | Nonancient | Yes | Yes | Yes | 71 | 71 |
| UD1 | Name unresolved in source mapping | Nonancient | Yes | Yes | Yes | 225 | 212 |
| UDP | Name unresolved in source mapping | Nonancient | Yes | Yes | Yes | 662 | 518 |
| UMP | Name unresolved in source mapping | Nonancient | Yes | Yes | Yes | 151 | 134 |
| UPG | Name unresolved in source mapping | Nonancient | Yes | Yes | Yes | 276 | 257 |
| URA | Uracil | Nonancient | Yes | Yes | Yes | 190 | 159 |
| URC | Name unresolved in source mapping | Nonancient | Yes | Yes | Yes | 127 | 109 |
| URE | Name unresolved in source mapping | Nonancient | Yes | Yes | Yes | 909 | 558 |
| URI | Name unresolved in source mapping | Nonancient | Yes | Yes | Yes | 79 | 79 |
| URO | Name unresolved in source mapping | Nonancient | Yes | Yes | Yes | 9 | 9 |
| UTP | Name unresolved in source mapping | Nonancient | Yes | Yes | Yes | 107 | 86 |
| VAL | Name unresolved in source mapping | Nonancient | Yes | Yes | Yes | 144 | 129 |
| VD3 | Name unresolved in source mapping | Nonancient | Yes | Yes | Yes | 186 | 152 |
| VDX | Calcitriol | Nonancient | Yes | Yes | Yes | 244 | 226 |
| VDY | Name unresolved in source mapping | Nonancient | Yes | Yes | Yes | 28 | 28 |
| XAN | Xanthine | Nonancient | Yes | Yes | Yes | 43 | 43 |
| XUL | Ribulose | Nonancient | Yes | Yes | Yes | 11 | 11 |
| YCM | Name unresolved in source mapping | Nonancient | Yes | Yes | Yes | 539 | 278 |

*Ancient status uses the prespecified ancient-metabolite list; 34 of those codes were present. Raw proteins are distinct reviewed human proteins assigned before feature-degree filtering. Cap-100 proteins are retained after restricting features to degree 2–100.*

**Supplementary Table S2A. Global topology of ancient and non-ancient retained features.**

| **Partition** | **Features** | **Nodes** | **Edges** | **Components** | **Largest nodes** | **Largest fraction** | **Top-two fraction** | **Edges/feature** |
| --- | --- | --- | --- | --- | --- | --- | --- | --- |
| ancient only | 504 | 1509 | 134801 | 2 | 1507 | 99.9% | 100.0% | 267.5 |
| non-ancient only | 15521 | 3586 | 739320 | 1 | 3586 | 100.0% | 100.0% | 47.6 |
| ancient involving | 7009 | 2976 | 552990 | 1 | 2976 | 100.0% | 100.0% | 78.9 |
| complete retained union | 22530 | 3786 | 951715 | 1 | 3786 | 100.0% | 100.0% | 42.2 |

*Features are single metabolites or exact attractive/repulsive COLIG states retained at degree 2–100. Mixed ancient–non-ancient pairs belong to ancient-involving but neither exclusive partition.*

**Supplementary Table S2B. Ancient and non-ancient pathway reconstruction by pathway class.**

| **Partition** | **Pathway class** | **Pathways** | **Protein coverage** | **Largest component** | **Top two components** | **One component** | **≤2 components** |
| --- | --- | --- | --- | --- | --- | --- | --- |
| ancient involving | Enzyme-only | 122 | 39.4% | 61.3% | 88.5% | 27 | 81 |
| ancient involving | Mixed | 1315 | 46.8% | 53.7% | 70.4% | 69 | 314 |
| ancient involving | Strict non-enzyme | 104 | 31.2% | 53.9% | 82.0% | 16 | 62 |
| ancient only | Enzyme-only | 122 | 19.7% | 46.2% | 78.3% | 9 | 61 |
| ancient only | Mixed | 1315 | 20.7% | 32.2% | 49.8% | 6 | 136 |
| ancient only | Strict non-enzyme | 104 | 13.9% | 42.0% | 73.8% | 6 | 51 |
| complete retained union | Enzyme-only | 122 | 54.0% | 69.4% | 92.7% | 41 | 96 |
| complete retained union | Mixed | 1315 | 63.1% | 66.8% | 82.2% | 209 | 553 |
| complete retained union | Strict non-enzyme | 104 | 45.1% | 63.3% | 88.3% | 29 | 72 |
| non-ancient only | Enzyme-only | 122 | 50.1% | 65.8% | 90.0% | 38 | 87 |
| non-ancient only | Mixed | 1315 | 57.2% | 61.2% | 77.2% | 145 | 422 |
| non-ancient only | Strict non-enzyme | 104 | 39.9% | 60.1% | 87.9% | 21 | 71 |

*One-component and ≤2-component counts use all eligible proteins, including isolated eligible proteins as components of size one. The ≤2 category includes one-component pathways.*

**Supplementary Table S2C. Exact bipartite degree-preserving null separates general connectivity from metabolite-specific placement.**

| **Network** | **Observed pathway edges** | **Null mean edges** | **Observed/null** | **P** | **Observed GCC** | **Observed GCC fraction** | **Null mean GCC** |
| --- | --- | --- | --- | --- | --- | --- | --- |
| Cap 100 | 698 | 640.1 | 1.090 | 0.005 | 3786 | 99.8% | 3782.9 |
| Uncapped | 1668 | 1678.5 | 0.994 | 0.995 | 5426 | 100.0% | 5425.9 |

*Exact bipartite double-edge swaps preserve every protein and feature degree while changing protein–feature assignments. GCC denotes the giant (largest) connected component. P values are empirical one-sided values from 199 retained null replicates.*

**Supplementary Table S3A. Structural-layer and feature-type network ablation audit.**

| **Layer or feature group** | **Retained features** | **Nodes** | **Projected edges** | **Edges beyond STRING** |
| --- | --- | --- | --- | --- |
| standalone monomer | 14011 | 2776 | 555649 | 554121 |
| dimer interface | 1456 | 1166 | 82147 | 81559 |
| dimer noninterface | 20059 | 2414 | 548970 | 547407 |
| single metabolite | 124 | 1809 | 112292 | 111740 |
| attractive COLIG | 2536 | 2266 | 140265 | 139591 |
| repulsive COLIG | 19870 | 3622 | 876791 | 874379 |

*Each group was rebuilt independently using the same feature-degree range of 2–100. Counts are descriptive and reflect both biological assignments and feature-degree distributions.*

**Supplementary Table S3B. Layer-specific pathway reconstruction and incremental coverage beyond STRING.**

| **Layer or feature group** | **Pathway class** | **n** | **Layer coverage** | **STRING + layer** | **Coverage gain** | **Union largest** | **Union top two** |
| --- | --- | --- | --- | --- | --- | --- | --- |
| standalone monomer | Strict non-enzyme | 104 | 33.9% | 79.0% | 8.9 pp | 84.1% | 94.7% |
| standalone monomer | Enzyme-only | 68 | 29.9% | 77.7% | 10.6 pp | 77.5% | 89.7% |
| standalone monomer | Mixed | 1283 | 44.7% | 85.3% | 8.2 pp | 83.7% | 93.5% |
| dimer interface | Strict non-enzyme | 104 | 13.7% | 72.0% | 1.8 pp | 76.3% | 90.1% |
| dimer interface | Enzyme-only | 68 | 20.4% | 68.5% | 1.4 pp | 65.2% | 83.4% |
| dimer interface | Mixed | 1283 | 21.3% | 79.3% | 2.2 pp | 75.6% | 88.0% |
| dimer noninterface | Strict non-enzyme | 104 | 22.5% | 75.8% | 5.6 pp | 79.7% | 92.5% |
| dimer noninterface | Enzyme-only | 68 | 50.3% | 81.0% | 13.9 pp | 79.8% | 93.2% |
| dimer noninterface | Mixed | 1283 | 44.1% | 84.8% | 7.7 pp | 83.9% | 93.2% |
| single metabolite | Strict non-enzyme | 104 | 11.3% | 74.2% | 4.0 pp | 77.5% | 91.0% |
| single metabolite | Enzyme-only | 68 | 27.1% | 75.2% | 8.1 pp | 73.6% | 89.0% |
| single metabolite | Mixed | 1283 | 21.3% | 80.0% | 2.9 pp | 76.8% | 88.8% |
| attractive COLIG | Strict non-enzyme | 104 | 20.4% | 77.2% | 7.0 pp | 80.2% | 92.4% |
| attractive COLIG | Enzyme-only | 68 | 23.0% | 74.6% | 7.5 pp | 74.7% | 87.6% |
| attractive COLIG | Mixed | 1283 | 25.3% | 81.6% | 4.5 pp | 78.0% | 89.8% |
| repulsive COLIG | Strict non-enzyme | 104 | 39.5% | 80.2% | 10.1 pp | 84.8% | 94.7% |
| repulsive COLIG | Enzyme-only | 68 | 61.2% | 87.3% | 20.2 pp | 84.8% | 96.3% |
| repulsive COLIG | Mixed | 1283 | 57.7% | 88.1% | 11.0 pp | 88.0% | 95.9% |

*Coverage gain is the percentage-point difference between STRING+layer and the corresponding STRING-only mean on the same fixed eligible cohort.*

**Supplementary Table S3C. Tissue-expression and shared-subcellular-location feasibility filters.**

| **Filter** | **Pathway class** | **n** | **Coverage** | **Largest** | **Top two** | **One component** | **≤2 components** |
| --- | --- | --- | --- | --- | --- | --- | --- |
| Tissue: cerebral cortex | Strict non-enzyme | 104 | 61.3% | 71.1% | 87.5% | 44 | 69 |
| Tissue: cerebral cortex | Enzyme-only | 68 | 69.8% | 72.7% | 87.5% | 30 | 43 |
| Tissue: cerebral cortex | Mixed | 1283 | 78.5% | 79.6% | 89.8% | 476 | 775 |
| Tissue: liver | Strict non-enzyme | 104 | 56.8% | 68.6% | 87.3% | 37 | 66 |
| Tissue: liver | Enzyme-only | 68 | 79.8% | 81.0% | 93.5% | 34 | 54 |
| Tissue: liver | Mixed | 1283 | 79.1% | 80.1% | 90.2% | 471 | 765 |
| Tissue: heart muscle | Strict non-enzyme | 104 | 58.7% | 69.3% | 87.1% | 39 | 64 |
| Tissue: heart muscle | Enzyme-only | 68 | 62.5% | 69.3% | 85.2% | 26 | 42 |
| Tissue: heart muscle | Mixed | 1283 | 79.2% | 80.2% | 90.3% | 475 | 787 |
| Tissue: lung | Strict non-enzyme | 104 | 59.5% | 70.2% | 87.7% | 41 | 66 |
| Tissue: lung | Enzyme-only | 68 | 68.0% | 71.7% | 86.8% | 27 | 43 |
| Tissue: lung | Mixed | 1283 | 81.1% | 81.7% | 91.3% | 523 | 818 |
| Tissue: colon | Strict non-enzyme | 104 | 55.2% | 67.3% | 86.5% | 36 | 66 |
| Tissue: colon | Enzyme-only | 68 | 71.7% | 74.9% | 89.3% | 28 | 43 |
| Tissue: colon | Mixed | 1283 | 80.7% | 81.4% | 91.0% | 525 | 818 |
| Tissue: kidney | Strict non-enzyme | 104 | 59.8% | 70.5% | 88.6% | 40 | 69 |
| Tissue: kidney | Enzyme-only | 68 | 70.9% | 74.5% | 91.7% | 25 | 48 |
| Tissue: kidney | Mixed | 1283 | 80.7% | 81.4% | 91.1% | 512 | 818 |
| Tissue: pancreas | Strict non-enzyme | 104 | 52.4% | 65.8% | 85.6% | 34 | 63 |
| Tissue: pancreas | Enzyme-only | 68 | 70.9% | 73.7% | 88.3% | 30 | 43 |
| Tissue: pancreas | Mixed | 1283 | 77.9% | 79.0% | 89.2% | 470 | 759 |
| Shared UniProt localization | Strict non-enzyme | 104 | 80.8% | 85.3% | 94.9% | 70 | 87 |
| Shared UniProt localization | Enzyme-only | 68 | 85.1% | 84.3% | 94.7% | 42 | 56 |
| Shared UniProt localization | Mixed | 1283 | 87.2% | 87.2% | 95.6% | 685 | 1026 |

*Tissue filters require Human Protein Atlas consensus expression in the named tissue. The localization filter requires at least one shared UniProt subcellular-location term. These are permissive feasibility screens, not cell-type-resolved occupancy measurements.*

**Supplementary Table S4A. Feature-degree cap sensitivity.**

| **Model** | **Pathway class** | **n** | **Coverage** | **Largest** | **Top two** | **One component** | **≤2 components** |
| --- | --- | --- | --- | --- | --- | --- | --- |
| cap 50 | Strict non-enzyme | 104 | 39.2% | 58.3% | 84.4% | 23 | 66 |
| cap 50 | Enzyme-only | 68 | 52.3% | 63.8% | 86.7% | 18 | 43 |
| cap 50 | Mixed | 1283 | 46.6% | 51.4% | 67.7% | 56 | 253 |
| STRING+cap 50 | Strict non-enzyme | 104 | 81.9% | 85.5% | 94.7% | 70 | 85 |
| STRING+cap 50 | Enzyme-only | 68 | 84.5% | 82.1% | 94.0% | 39 | 54 |
| STRING+cap 50 | Mixed | 1283 | 85.6% | 84.5% | 94.0% | 634 | 938 |
| cap 75 | Strict non-enzyme | 104 | 41.8% | 60.4% | 85.4% | 26 | 68 |
| cap 75 | Enzyme-only | 68 | 58.5% | 68.0% | 89.2% | 20 | 43 |
| cap 75 | Mixed | 1283 | 55.4% | 59.3% | 74.8% | 124 | 370 |
| STRING+cap 75 | Strict non-enzyme | 104 | 82.3% | 86.2% | 95.1% | 71 | 87 |
| STRING+cap 75 | Enzyme-only | 68 | 86.2% | 83.8% | 95.4% | 40 | 55 |
| STRING+cap 75 | Mixed | 1283 | 87.5% | 87.0% | 95.5% | 706 | 1016 |
| cap 100 | Strict non-enzyme | 104 | 44.6% | 62.4% | 86.9% | 29 | 69 |
| cap 100 | Enzyme-only | 68 | 61.9% | 71.1% | 91.0% | 23 | 50 |
| cap 100 | Mixed | 1283 | 61.3% | 64.6% | 79.4% | 170 | 459 |
| STRING+cap 100 | Strict non-enzyme | 104 | 82.5% | 86.3% | 95.1% | 73 | 87 |
| STRING+cap 100 | Enzyme-only | 68 | 88.0% | 86.1% | 96.4% | 44 | 59 |
| STRING+cap 100 | Mixed | 1283 | 88.9% | 88.8% | 96.4% | 767 | 1068 |
| cap 125 | Strict non-enzyme | 104 | 46.7% | 64.2% | 88.4% | 30 | 74 |
| cap 125 | Enzyme-only | 68 | 66.9% | 73.8% | 92.3% | 26 | 51 |
| cap 125 | Mixed | 1283 | 65.3% | 68.3% | 82.4% | 222 | 527 |
| STRING+cap 125 | Strict non-enzyme | 104 | 82.7% | 87.3% | 95.6% | 74 | 89 |
| STRING+cap 125 | Enzyme-only | 68 | 89.2% | 86.8% | 97.2% | 45 | 60 |
| STRING+cap 125 | Mixed | 1283 | 89.8% | 89.9% | 96.8% | 819 | 1092 |
| cap 150 | Strict non-enzyme | 104 | 50.8% | 66.8% | 90.0% | 33 | 77 |
| cap 150 | Enzyme-only | 68 | 74.2% | 79.3% | 94.6% | 32 | 55 |
| cap 150 | Mixed | 1283 | 68.7% | 71.7% | 85.5% | 261 | 599 |
| STRING+cap 150 | Strict non-enzyme | 104 | 83.4% | 88.2% | 96.5% | 74 | 93 |
| STRING+cap 150 | Enzyme-only | 68 | 92.0% | 90.1% | 97.9% | 49 | 61 |
| STRING+cap 150 | Mixed | 1283 | 90.6% | 91.0% | 97.3% | 853 | 1113 |
| cap 250 | Strict non-enzyme | 104 | 59.2% | 74.3% | 94.0% | 43 | 84 |
| cap 250 | Enzyme-only | 68 | 83.0% | 84.9% | 96.7% | 41 | 59 |
| cap 250 | Mixed | 1283 | 77.0% | 79.8% | 91.2% | 419 | 792 |
| STRING+cap 250 | Strict non-enzyme | 104 | 86.3% | 90.4% | 97.4% | 79 | 95 |
| STRING+cap 250 | Enzyme-only | 68 | 93.7% | 91.6% | 98.7% | 51 | 63 |
| STRING+cap 250 | Mixed | 1283 | 93.0% | 93.4% | 98.4% | 954 | 1173 |
| uncapped | Strict non-enzyme | 104 | 99.8% | 99.8% | 100.0% | 103 | 104 |
| uncapped | Enzyme-only | 68 | 100.0% | 100.0% | 100.0% | 68 | 68 |
| uncapped | Mixed | 1283 | 100.0% | 100.0% | 100.0% | 1281 | 1283 |
| STRING+uncapped | Strict non-enzyme | 104 | 100.0% | 100.0% | 100.0% | 104 | 104 |
| STRING+uncapped | Enzyme-only | 68 | 100.0% | 100.0% | 100.0% | 68 | 68 |
| STRING+uncapped | Mixed | 1283 | 100.0% | 100.0% | 100.0% | 1283 | 1283 |

*Caps apply to feature degree, not final protein-network degree. Uncapped retains every feature assigned to at least two proteins.*

**Supplementary Table S4B. Pathway-size-stratified reconstruction sensitivity.**

| **Model** | **Pathway class** | **Eligible size bin** | **n** | **Coverage** | **Largest** | **Top two** |
| --- | --- | --- | --- | --- | --- | --- |
| STRING | Enzyme-only | 2-10 | 64 | 68.0% | 67.7% | 85.3% |
| STRING | Enzyme-only | 11-30 | 4 | 52.2% | 25.4% | 41.5% |
| STRING | Mixed | 2-10 | 803 | 73.6% | 71.5% | 87.3% |
| STRING | Mixed | 11-30 | 418 | 83.6% | 70.5% | 81.7% |
| STRING | Mixed | 31+ | 62 | 78.9% | 68.4% | 75.1% |
| STRING | Strict non-enzyme | 2-10 | 99 | 69.6% | 74.9% | 89.9% |
| STRING | Strict non-enzyme | 11-30 | 4 | 83.3% | 76.4% | 82.6% |
| STRING | Strict non-enzyme | 31+ | 1 | 75.0% | 75.0% | 77.5% |
| LIGMAP | Enzyme-only | 2-10 | 64 | 59.8% | 69.5% | 90.5% |
| LIGMAP | Enzyme-only | 11-30 | 4 | 96.0% | 96.0% | 100.0% |
| LIGMAP | Mixed | 2-10 | 803 | 51.8% | 59.3% | 78.9% |
| LIGMAP | Mixed | 11-30 | 418 | 75.4% | 71.3% | 78.8% |
| LIGMAP | Mixed | 31+ | 62 | 88.1% | 87.5% | 90.2% |
| LIGMAP | Strict non-enzyme | 2-10 | 99 | 44.1% | 63.2% | 88.5% |
| LIGMAP | Strict non-enzyme | 11-30 | 4 | 41.5% | 33.2% | 42.1% |
| LIGMAP | Strict non-enzyme | 31+ | 1 | 100.0% | 100.0% | 100.0% |
| STRING+LIGMAP | Enzyme-only | 2-10 | 64 | 87.4% | 85.3% | 96.1% |
| STRING+LIGMAP | Enzyme-only | 11-30 | 4 | 98.1% | 98.1% | 100.0% |
| STRING+LIGMAP | Mixed | 2-10 | 803 | 84.6% | 85.3% | 95.5% |
| STRING+LIGMAP | Mixed | 11-30 | 418 | 95.9% | 94.4% | 97.8% |
| STRING+LIGMAP | Mixed | 31+ | 62 | 96.9% | 96.8% | 98.2% |
| STRING+LIGMAP | Strict non-enzyme | 2-10 | 99 | 82.2% | 86.6% | 95.5% |
| STRING+LIGMAP | Strict non-enzyme | 11-30 | 4 | 84.7% | 76.4% | 84.0% |
| STRING+LIGMAP | Strict non-enzyme | 31+ | 1 | 100.0% | 100.0% | 100.0% |
| FULL | Enzyme-only | 2-10 | 64 | 88.5% | 86.4% | 96.4% |
| FULL | Enzyme-only | 11-30 | 4 | 98.1% | 98.1% | 100.0% |
| FULL | Mixed | 2-10 | 803 | 85.1% | 85.6% | 95.8% |
| FULL | Mixed | 11-30 | 418 | 96.2% | 94.8% | 97.9% |
| FULL | Mixed | 31+ | 62 | 97.2% | 97.0% | 98.4% |
| FULL | Strict non-enzyme | 2-10 | 99 | 82.4% | 86.8% | 95.6% |
| FULL | Strict non-enzyme | 11-30 | 4 | 84.7% | 76.4% | 84.0% |
| FULL | Strict non-enzyme | 31+ | 1 | 100.0% | 100.0% | 100.0% |

*Size bins use the number of eligible proteins in the fixed comparison cohort.*

**Supplementary Table S4C. WikiPathways identifier-mapping and Reactome-overlap sensitivity.**

| **Mapping route** | **Mapped memberships** | **Model** | **Overlap band** | **n** | **Coverage** | **Largest** | **Top two** |
| --- | --- | --- | --- | --- | --- | --- | --- |
| STRING alias | 48.2% | STRING | all | 935 | 57.4% | 49.1% | 64.7% |
| STRING alias | 48.2% | STRING | Reactome overlap ge 0.5 | 62 | 64.1% | 62.3% | 77.2% |
| STRING alias | 48.2% | STRING | Reactome overlap lt 0.5 | 873 | 56.9% | 48.1% | 63.8% |
| STRING alias | 48.2% | LIGMAP | all | 935 | 73.3% | 75.7% | 86.0% |
| STRING alias | 48.2% | LIGMAP | Reactome overlap ge 0.5 | 62 | 59.6% | 62.6% | 78.3% |
| STRING alias | 48.2% | LIGMAP | Reactome overlap lt 0.5 | 873 | 74.3% | 76.6% | 86.6% |
| STRING alias | 48.2% | STRING+LIGMAP | all | 935 | 86.1% | 87.0% | 94.8% |
| STRING alias | 48.2% | STRING+LIGMAP | Reactome overlap ge 0.5 | 62 | 82.2% | 83.4% | 93.0% |
| STRING alias | 48.2% | STRING+LIGMAP | Reactome overlap lt 0.5 | 873 | 86.4% | 87.2% | 94.9% |
| STRING alias | 48.2% | FULL | all | 935 | 87.2% | 87.9% | 95.5% |
| STRING alias | 48.2% | FULL | Reactome overlap ge 0.5 | 62 | 84.1% | 84.2% | 94.1% |
| STRING alias | 48.2% | FULL | Reactome overlap lt 0.5 | 873 | 87.5% | 88.1% | 95.6% |
| NCBI symbol | 47.9% | STRING | all | 935 | 56.5% | 48.4% | 64.1% |
| NCBI symbol | 47.9% | STRING | Reactome overlap ge 0.5 | 62 | 63.3% | 59.6% | 75.9% |
| NCBI symbol | 47.9% | STRING | Reactome overlap lt 0.5 | 873 | 56.1% | 47.6% | 63.2% |
| NCBI symbol | 47.9% | LIGMAP | all | 935 | 73.2% | 75.5% | 85.9% |
| NCBI symbol | 47.9% | LIGMAP | Reactome overlap ge 0.5 | 62 | 60.9% | 63.6% | 79.1% |
| NCBI symbol | 47.9% | LIGMAP | Reactome overlap lt 0.5 | 873 | 74.1% | 76.4% | 86.3% |
| NCBI symbol | 47.9% | STRING+LIGMAP | all | 935 | 85.8% | 86.6% | 94.5% |
| NCBI symbol | 47.9% | STRING+LIGMAP | Reactome overlap ge 0.5 | 62 | 82.1% | 83.3% | 92.9% |
| NCBI symbol | 47.9% | STRING+LIGMAP | Reactome overlap lt 0.5 | 873 | 86.0% | 86.8% | 94.6% |
| NCBI symbol | 47.9% | FULL | all | 935 | 86.9% | 87.5% | 95.2% |
| NCBI symbol | 47.9% | FULL | Reactome overlap ge 0.5 | 62 | 84.1% | 84.1% | 94.1% |
| NCBI symbol | 47.9% | FULL | Reactome overlap lt 0.5 | 873 | 87.1% | 87.7% | 95.3% |

*Overlap bands compare each WikiPathways membership with its best Reactome overlap. The two mapping routes were evaluated independently.*

**Supplementary Table S4D. Exact BioLiP2 human protein–metabolite contact recall by LIGMAP scope.**

| **LIGMAP scope** | **Experimental contacts** | **Recovered** | **Recall** |
| --- | --- | --- | --- |
| all single metabolite raw | 2314 | 849 | 36.7% |
| dimer interface all | 2314 | 65 | 2.8% |
| dimer interface att | 2314 | 0 | 0.0% |
| dimer interface rep | 2314 | 0 | 0.0% |
| dimer monomer all | 2314 | 529 | 22.9% |
| dimer monomer att | 2314 | 0 | 0.0% |
| dimer monomer rep | 2314 | 0 | 0.0% |
| standalone monomer all | 2314 | 561 | 24.2% |
| standalone monomer att | 2314 | 0 | 0.0% |
| standalone monomer rep | 2314 | 0 | 0.0% |
| cap 50 | 2314 | 14 | 0.6% |
| cap 75 | 2314 | 27 | 1.2% |
| cap 100 | 2314 | 28 | 1.2% |
| cap 125 | 2314 | 38 | 1.6% |
| cap 150 | 2314 | 47 | 2.0% |
| cap 250 | 2314 | 66 | 2.9% |
| cap none | 2314 | 849 | 36.7% |

*Raw single-metabolite incidence is the appropriate object for contact recall. Cap-100 is a topology-selective projection and excludes broadly assigned features.*

**Supplementary Table S4E. Within-pathway reference-pair recovery by endpoint-degree quartile.**

| **Model** | **Maximum endpoint-degree quartile** | **Reference pairs** | **Recovered pairs** | **Recovery** |
| --- | --- | --- | --- | --- |
| STRING | 1 | 4061 | 229 | 5.6% |
| STRING | 2 | 14907 | 1100 | 7.4% |
| STRING | 3 | 27725 | 1570 | 5.7% |
| STRING | 4 | 57767 | 3227 | 5.6% |
| LIGMAP | 1 | 4061 | 20 | 0.5% |
| LIGMAP | 2 | 14907 | 231 | 1.5% |
| LIGMAP | 3 | 27725 | 1695 | 6.1% |
| LIGMAP | 4 | 57767 | 20293 | 35.1% |
| STRING+LIGMAP | 1 | 4061 | 244 | 6.0% |
| STRING+LIGMAP | 2 | 14907 | 1292 | 8.7% |
| STRING+LIGMAP | 3 | 27725 | 3099 | 11.2% |
| STRING+LIGMAP | 4 | 57767 | 22101 | 38.3% |
| FULL | 1 | 4061 | 274 | 6.7% |
| FULL | 2 | 14907 | 1374 | 9.2% |
| FULL | 3 | 27725 | 3307 | 11.9% |
| FULL | 4 | 57767 | 22371 | 38.7% |

*Pairs are stratified by the larger endpoint degree. The strong increase in LIGMAP recovery across quartiles documents hub concentration.*
